# Protein-lipid charge interactions control the folding of OMPs into asymmetric membranes

**DOI:** 10.1101/2023.04.20.537663

**Authors:** Jonathan Machin, Antreas C. Kali, Neil A. Ranson, Sheena E. Radford

**Affiliations:** Astbury Centre for Structural Molecular Biology, School of Molecular and Cellular Biology, Faculty of Biological Sciences, University of Leeds, LS2 9JT, United Kingdom; Astbury Centre for Structural Molecular Biology and Leeds Institute of Cardiovascular and Metabolic Medicine, School of Medicine, University of Leeds, LS2 9JT, United Kingdom

## Abstract

Biological membranes consist of two leaflets of phospholipid molecules that form a bilayer, and typically the composition of lipids in each leaflet is distinct. This asymmetry is created and maintained *in vivo* by dedicated biochemical pathways, but difficulties in creating stable asymmetric membranes *in vitro* have restricted our understanding of how bilayer asymmetry modulates the folding, stability and function of membrane proteins. Here we employ cyclodextrin mediated lipid exchange to generate asymmetric liposomes and use these to characterize the stability and folding kinetics of two bacterial outer membrane proteins (OMPs). We show that excess negative charge in the outer leaflet of a liposome impedes the membrane insertion and folding of OmpA and BamA, while excess negative charge in the inner leaflet accelerates their folding, relative to symmetric liposomes with the same membrane composition. Three positively charged residues in the extracellular loops of OmpA that play a critical role in folding are identified using molecular dynamics simulations and mutational analyses. Bioinformatic analysis was then used to identify a conserved patch of positive residues in the extracellular loops of OMPs generally that lies 6-8Å from the membrane surface. Together, the the results rationalise the well known ‘positive outside’ rule for OMP sequences and suggest new insights into the mechanisms that drive OMP folding and assembly *in vitro* and *in vivo*.

## Introduction

Membrane proteins are ubiquitous, carrying out many essential functions in biology, and are therefore major drug targets ^1, 2^. Recent progress has been made in understanding how membrane proteins fold ^3^, but while lipid-protein interactions are important, their precise roles remain unclear, with most information gleaned empirically on a case-by-case basis ^4, 5^. Most biological membranes have an asymmetric lipid composition in each leaflet of their bilayer. This asymmetry is membrane-type and cell-status dependent ^6^, and the plethora of enzymes dedicated to creating and maintaining bilayer asymmetry ^7, 8^, as well as disease states featuring mis-regulated asymmetry ^9^, demonstrate its importance. However, generating stable, lipid asymmetric systems of the quality and quantity needed for *in vitro* folding studies is challenging, so little is currently known about how membrane asymmetry and protein folding interplay.

Methods to generate asymmetric phospholipid bilayers do exist, including supported bilayers ^10, 11^, phase-transfer approaches ^12, 13^ and liposome hemifusion ^14^. Asymmetry can also be generated by cyclodextrin (CD)-mediated lipid exchange ^15, 16^ which has been used to generate liposomes with asymmetric bilayers ^17, 18^. In a few cases, these have been used to study membrane protein folding, with the rate of folding of perfringolysin O ^19^ and the ‘pH low insertion peptide’ (pHLIP) ^20^ each being modulated by charge asymmetries across the bilayer. However, both perfringolysin O and pHLIP have stable, water-soluble forms, and only insert into membranes under specific conditions ^21, 22^. It is thus hard to generalise these finding to integral membrane proteins, which require the membrane to adopt their native fold.

Outer membrane proteins (OMPs) from Gram-negative bacteria ^23, 24^ have a trans-membrane β- barrel fold, where membrane-spanning β-strands are linked by long extracellular loops and short intracellular turns ^25^. *In vitro* folding studies of OMPs of different sequence and size have shown that the membrane plays an important role in regulating folding ^26^. For example, folding is faster when bilayers contain lipids with shorter acyl-chains ^27^, less saturated lipids ^28^, or more membrane defects ^29^, all of which facilitate the OMP partitioning into the membrane. Lipid head groups also modulate folding, with phosphoethanolamine (PE) and phosphoglycerol (PG) introducing a kinetic barrier for folding in C_10:0_ lipids ^30, 31^. However, recent work with C_14:0_ lipids did not show this kinetic barrier, perhaps due to the additional kinetic barrier of a thicker membrane dominating folding ^32^. The primary sequence of an OMP is also critical, and may be more significant than the membrane properties ^32^, a concept supported by mutational analysis of the folding efficiency of OmpA, EspP and OmpC variants *in vivo* ^33^. While the folding of OMPs into membranes has been well studied over decades of research (reviewed in ^23, 34^), the role of membrane asymmetry in OMP folding has not been explored in detail to date.

Here, we used CD-mediated lipid exchange to generate charge-asymmetric liposomes, using DMPC, DMPG, DMPE and DMPS lipids, and validate their asymmetry using measurements and predictions of their ζ-potential. We show for two model OMPs, the 8-stranded OmpA and the 16- stranded BamA, that folding rates and stability are modulated by a leaflet-specific distribution of negatively charged lipid head groups. Using molecular dynamics (MD) simulations, we identify specific, positively charged residues in the extracellular loops of OmpA that interact with lipids and show that they are critical for OmpA folding *in vitro*. Bioinformatics analysis of >300 structures and >19,000 sequences of OMPs revealed a highly-conserved enrichment of positively-charged residues in the extracellular loops close to the membrane surface. Collectively, these results reveal that efficient OMP folding requires a previously-uncharted synergy between the lipid charge in each leaflet of a bilayer and a signature region of charges in the extracellular loops of the folding OMP. They also offer insights into the molecular mechanisms that drive OMP folding and stability into asymmetric membranes *in vivo,* and suggest how these can be controlled for biotechnological applications.

## Results

### Generating asymmetric liposomes

Charge distribution in membrane proteins is used to control protein topology and stability *in vivo* ^35^, such as the ‘positive-inside’ rule which modulates the orientation of proteins in the plasma membrane/bacterial inner membrane ^36, 37^. By contrast, the ‘positive-outside’ rule for OMPs, with more Lys/Arg residues in the extracellular loops than the intracellular turns, is postulated to stabilize OMPs via their interaction with lipopolysaccharide (LPS) in the highly asymmetric bacterial OM ^38, 39^. However, shortening the extracellular loops of OMPs either individually or in combination, thus removing positive charge, does not alter their folding topology, suggesting that charge has a different, currently unknown, role in OMP assembly ^40, 41^.

To determine the effect(s) of lipid charge asymmetry on OMP folding, membrane systems based on dimyristoyl-phosphatidylcholine and -phosphoglycerol (DMPC and DMPG) were created. These lipids have the same C_14:0_ acyl-chains, similar T_m_s (∼24 and 23 °C, respectively (**Extended Data Fig. 1**)) and similar head group sizes, but different charge: DMPC is a neutral zwitterion while DMPG is negatively charged (**Fig. 1a**). DMPC-DMPG asymmetric liposomes were generated by methyl beta-cyclodextrin (MβCD)-mediated exchange (**Fig. 1b**). Note that hereinafter, symmetric and asymmetric lipid mixes are indicated by an s- and a- prefix respectively, and asymmetric liposomes are denoted as donor-lipid/acceptor-liposome; thus DMPG/PC indicates DMPG lipids exchanged into the outer leaflet of DMPC liposomes. (All lipid ratios are *mol/mol* unless otherwise indicated). Following lipid exchange, liposome integrity and size was confirmed using cryoEM and dynamic light scattering (DLS) (**Fig. 1c-d**). Efficient removal of MβCD from the final liposome preparation was confirmed via detection of residual sugar (Methods) (**Fig. S1**). The ability of MβCD to mediate exchange between DMPC and DMPG lipids was confirmed using a fluorescent marker (DPPE-Rhodamine) to indicate exchange (**Extended Data Fig. 2** and Methods). Thin layer chromatography (TLC), quantified by densitometry, was then used to measure label-free lipid exchange. Controls confirmed that DMPG and DMPC staining intensity was directly proportional to the amount of lipid loaded (within an error of <3%) (**Fig. S2**), allowing the DMPC:DMPG ratio to be determined and hence the extent of lipid exchange to be quantified (for example, **Fig. 1e**). Label-free DMPC-DMPG liposome asymmetry was then confirmed by determining the ζ-potential, a measure of particle surface charge, allowing quantitation of the amount of neutral DMPC and the negatively-charged DMPG in the solvent-exposed outer leaflet of liposomes (**Fig. S3**). Together with the total lipid ratio, ζ-potential thus provides a direct readout of lipid asymmetry (**Fig. 1f**). For example, the green circled a-DMPG/PC exchange sample has a total DMPG fraction of ∼25 % (lower x-axis) and a ζ-potential -26 mV. These data fall onto the dashed, theoretical “asymmetry” line, confirming this liposome is asymmetric, and allowing the outer leaflet DMPG content to be read from the upper x-axis. The asymmetric liposomes generated were stable for at least 72 h and in the presence of 8 M urea (**Fig. S3**).

**Figure 1:**
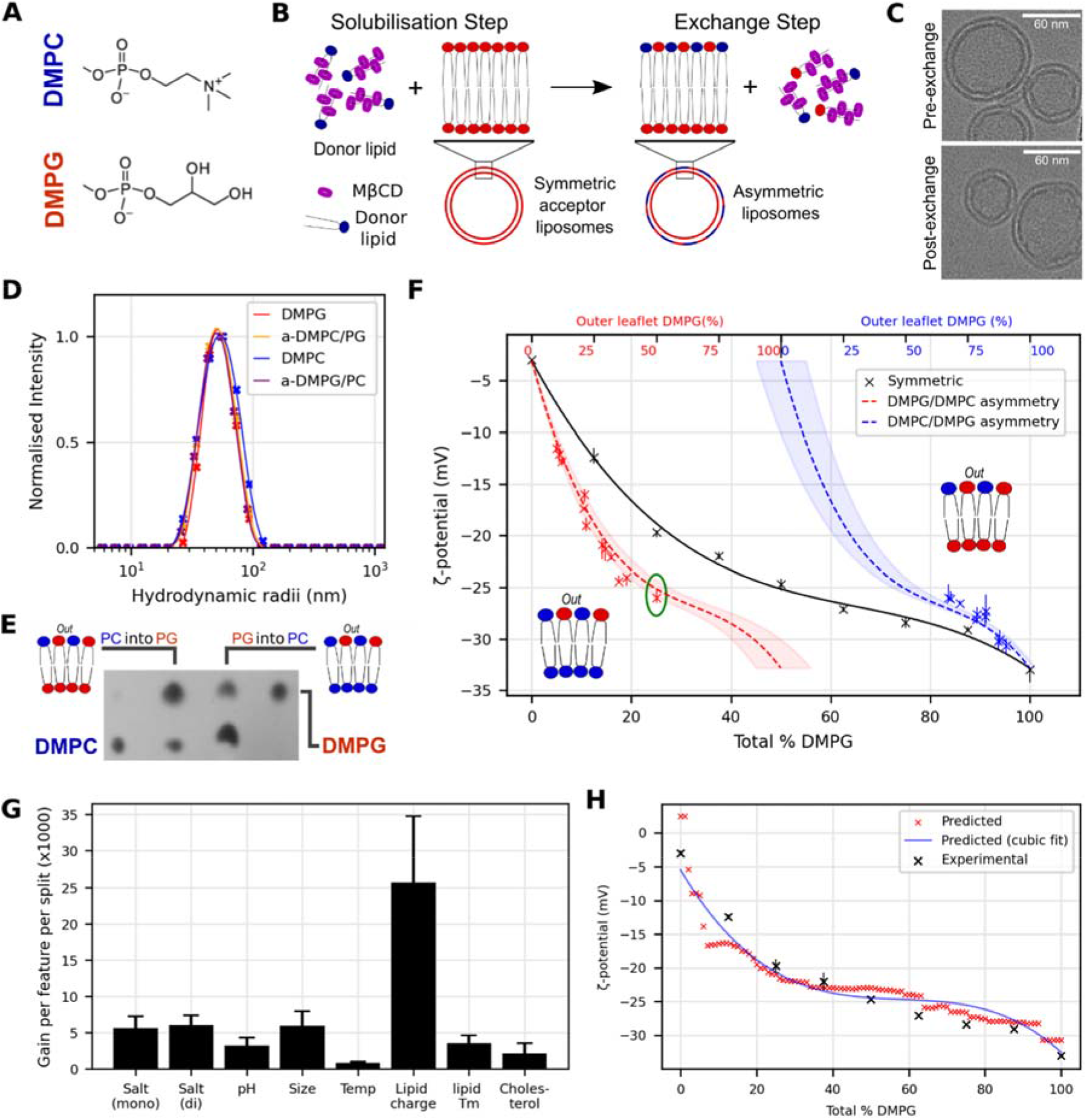
Generating and validating asymmetric LUVs. **(A)** Headgroup structure of DMPG and DMPC lipids (red and blue respectively, colours used throughout). **(B)** Overview of asymmetric liposome generation by MβCD-mediated exchange. **(C)** Pre-/post-exchange liposomes imaged by cryo-EM, these are smaller than the DLS as small liposomes preferentially go into the ice. **(D)** Pre-/post-exchange liposome size by DLS. **(E)** Sample TLC plate showing the lipid content of DMPC and DMPG exchanged liposomes, as indicated. **(F)** ζ- potential by lipid content for symmetric (black line) and asymmetric liposomes (blue curve: DMPC/PG, red curve: DMPG/PC). Theoretical asymmetry lines are shown with 10% error margin. Generated asymmetric liposome samples (DMPC/PG: red, DMPG/PC: blue) shown with range bars from repeat ζ-potential measurements. Green circled measurement discussed in text. **(G)** Feature importance (gain per feature per split) in the liposome ζ-potential model, error bars are standard deviation of trained model ensemble (n=50). **(H)** Agreement between predicted (red) and experimental (black) values for DMPC-PG LUVs in the buffer conditions used here (100 mM NaCl, 20 mM Tris-Cl, pH 8.5).

### A predictive model for liposome ζ-potential

To improve our ability to define lipid asymmetry of different lipid compositions using experimental measurement of the ζ-potential, a machine learning model was constructed to predict ζ-potential (Methods). 315 data points were collated from the literature and the work presented here ^17, 42–54^. Lipid composition was simplified by parametrisation to: (1) average overall charge per lipid, (2) average T_m_ of all lipids, and (3) fraction of cholesterol present, and when combined with five additional liposome/buffer features yielded an optimised model with an average mean absolute error (MAE) ∼3.0 mV. Overall lipid charge dominates the model (**Fig. 1g**), and parameter ablation indicates that lipid charge, lipid T_m_ and salt concentration are the most predictive features (**Fig. S4**). The model prediction for DMPG and DMPC lipid mixtures (excluding measured data) aligns well with experimental data (MAE=0.86 mV, similar to the average experimental measurement range of 0.88 mV) (**Fig. 1h**). DMPC and DMPG lipids are well represented in the training data, but the ζ-potential trend of less well-represented lipids and their mixtures, such as DMPS-DMPC and DMPE-DMPG was also correctly predicted over the regions experimentally validated, but with a larger error (**Fig. S5**). The predicted and measured ζ-potentials, combined with our experimental data, confirm that stable asymmetric liposomes of up to ∼30% a-DMPC/PG and ∼50% a-DMPG/PC in the outer leaflet of the LUVs can be generated (**Fig. 1f**).

### Lipid asymmetry modulates the folding rate of an OMP

The effect of lipid asymmetry on OMP folding was next assessed by measuring the folding rate and urea-stability of OmpA folded into symmetric and asymmetric bilayers, using tryptophan fluorescence and cold SDS-PAGE (Methods). OmpA is a well-studied model for OMP folding *in vitro* ^55–57^, and contains two domains: an eight-stranded transmembrane β-barrel and a C-terminal (natively periplasmic) water soluble domain. This C-terminal domain cannot cross the bilayer, ensuring unidirectional membrane insertion ^58^ (confirmed via trypsin cleavage, **Extended Data Fig. 3**), but its presence has minimal effect on the folding kinetics of the transmembrane region^55^. This unidirectionality allows the effects of lipid asymmetry on the observed rate of OmpA folding and stability to urea denaturation to be determined. Folding kinetics were measured at 30°C to ensure that all membranes are in the fluid lipid phase, regardless of their composition (**Extended Data Fig. 1**), and fitted to a single exponential (Methods) to derive the observed rate constant of folding (**Extended Data Fig. 4**). As expected ^31^, OmpA folds efficiently (folding yield 79%) (**Fig. 2c**) into symmetric DMPC liposomes with a rate constant (*k_obs_*) of ∼0.5x10^-3^ s^-1^ (**Fig. 2a**). Addition of small amounts (10%) of DMPG into both leaflets (i.e. maintaining symmetry) slows folding slightly (40% decrease in *k_obs_*), while higher concentrations of symmetric DMPG accelerate folding substantially (e.g. a ∼5-fold increase in *k_obs_* is observed with 40 % DMPG) (**Fig. 2a**). Strikingly different results were observed for asymmetric membranes with the same outer leaflet lipid composition. For example, in liposomes containing ≥ 20 % DMPG in the outer leaflet and pure DMPC in the inner leaflet, OmpA failed to fold within the 15 h timeframe of the experiment (final urea concentration of 0.48 M) (**Fig. 2a**). By contrast, while OmpA folds more than 40-times more rapidly into symmetric membranes of pure DMPG compared with pure DMPC (compare **Figs. 2a-b**), titrating DMPC into the outer leaflet of DMPG liposomes increases the rate constant for folding ∼ 2-fold relative to symmetric liposomes with equivalent outer leaflet composition at all percent lipid compositions measured (**Fig. 2b**). The lipid composition in each leaflet of the bilayer thus has a dramatic effect on the rate of OmpA folding. Given the similarity in T_m_, area-per-lipid, and acyl-chain length of DMPC and DMPG, these effects presumably arise from the different lipid head group charge in the membrane leaflets.

**Figure 2:**
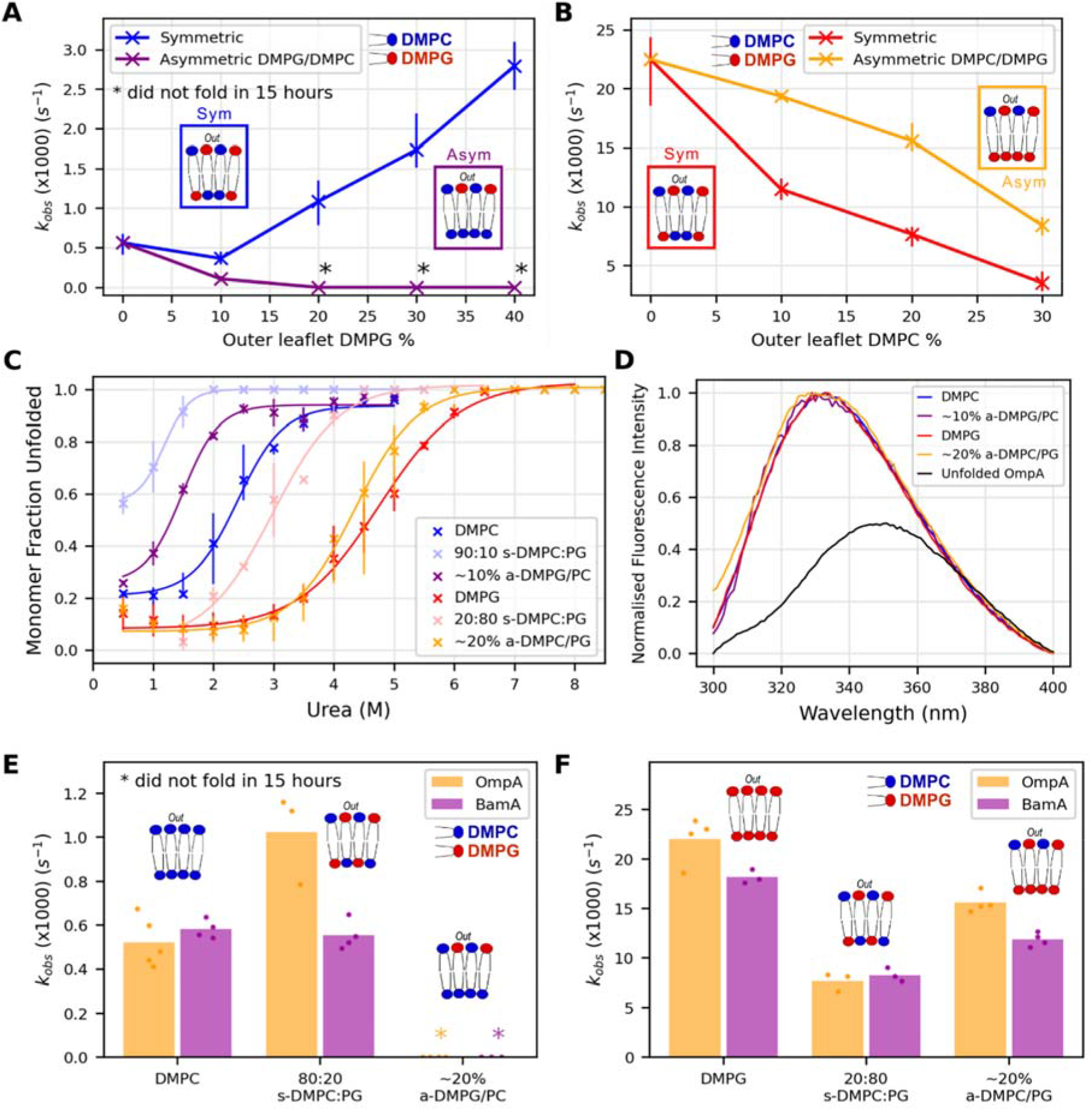
DMPC-DMPG lipid asymmetry significantly effects OMP folding rates. Folding rate constants (s^-1^) of OmpA into **(A)** a-DMPG/PC and **(B)** a-DMPC/PG asymmetric liposomes compared to symmetric liposomes with the same outer leaflet composition. Bars represent data ranges, * indicates the folding had not finished in 15 hours (< 75% folded). **(C)** Urea dependence of OmpA folding into DMPC-DMPG symmetric and asymmetric liposomes, as indicated. The lines are the fit to the average. **(D)** Tryptophan fluorescence emission spectra of OmpA folded into LUVs of different composition show that the protein does not unfold after overnight incubation at 30 °C in 7.5 M urea in any liposome. The spectrum of OmpA unfolded in 7.5 M urea in the absence of lipid is shown for comparison. **(E,F)** Observed folding rate constant (s^-1^) of OmpA/BamA into asymmetric and symmetric liposomes, demonstrating similar trends for the two proteins in each liposome type. * Indicates the folding was not complete (< 75% folded) in 15 hours and hence a rate constant could not be determined (Methods).

The stability of OmpA folded into symmetric and asymmetric bilayers was also assessed using cold SDS-PAGE (Methods). This assay makes use of the resistance of natively folded OmpA to denaturation by SDS in the absence of heating, enabling the fraction of folded/unfolded OmpA versus urea concentration to be determined using densitometry of bands in the gel (Methods) (**Fig 2c**, **Fig. S6**). OmpA has a higher apparent stability in symmetric DMPG liposomes compared with symmetric DMPC liposomes (urea concentration at the mid-point of each titration curve (P_m_) of 4.5 M and 2.3 M urea, respectively) (**Fig. 2c**). Unfolding of OmpA shows strong hysteresis, with the membrane embedded native protein being resilient to unfolding in all liposomes even in 8 M urea (**Fig. 2d**, **Fig. S7**) (hence equilibrium ΔG°_(eq)_ values could not be determined, consistent with previous reports ^59^). While addition of small amounts (20%) DMPC into the outer leaflet of DMPG LUVs has little effect on P_m_ (4.5 M urea), adding 20% DMPC symmetrically into both leaflets of the bilayer destabilised the protein (P_m_ 3 M urea) compared with pure DMPG LUVs. Adding 10 % DMPG asymmetrically into the outer leaflet of DMPC LUVs also destabilised the protein relative to pure DMPC liposomes (P_m_ 1.3 M urea), with a symmetric organisation of the same lipid composition having a greater effect (P_m_ ∼0.8 M urea) (**Fig. 2c**). These data were independently confirmed by assessing folding using tryptophan fluorescence (**Fig. S8**). Collectively, these results show that membrane asymmetry modulates both the rate and apparent stability of OmpA folding: an excess of DMPG (i.e. an excess of negative charge) in the outer leaflet slows folding and decreases P_m_, while an excess of DMPG in the inner leaflet enhances the rate of folding and increases P_m_ compared with symmetric liposomes with the same outer leaflet lipid composition.

To determine whether these effects are unique to OmpA, folding of the 16-stranded OMP BamA was also studied. Like OmpA, BamA has a large (47 kDa) water soluble domain that ensures the unidirectional folding of its 43 kDa transmembrane β-barrel domain (**Extended Data Fig. 3**). The folding kinetics of BamA into symmetric and asymmetric liposomes showed similar trends to OmpA: DMPG/PC asymmetry slows (or abrogates) folding while DMPC/PG asymmetry accelerates folding, relative to symmetric systems with the same outer leaflet composition (**Fig. 2e-f**, **Extended Data Fig. 4**). The trends observed for OmpA/BamA may therefore be general to OMPs, and suggests that lipid asymmetry, presumably the precise organisation of charge on each leaflet of the bilayer, plays a critical role in the folding rate and stability of OMP insertion into a membrane.

### Changes in OMP folding and stability arise from charge effects

To confirm whether the effects of lipid asymmetry on OMP folding are charge-mediated, or specific to DMPG and DMPC, we studied OmpA folding into membranes containing DMPS and DMPE (**Fig. 3a**). Like DMPG, DMPS has a net negative charge, while DMPE, like DMPC, is net neutral. As DMPS and DMPE have T_m_ values of 35 °C and 50 °C, respectively ^60^, to ensure all assays were performed using lipids in a fluid lipid phase DMPE and DMPS were used only at low concentrations (< 20%) with DMPG or DMPC. Lipid phase for the different lipid compositions was confirmed using a laurdan-based lipid-phase assay ^61^ (**Extended Data Fig. 1**). Asymmetric DMPS/PC and DMPE/PG LUVs were prepared, and validated by ζ-potential, TLC and DLS (**Extended Data Fig. 5**-**6**). The kinetics of OmpA folding into a-DMPS/PC showed that addition of DMPS into the outer leaflet of DMPC liposomes retards folding, akin to a-DMPG/PC (**Fig. 3b**). Unlike DMPG-PC lipid mixes, the stability of inserted OmpA is similar in the DMPS-PC symmetric and asymmetric membranes (**Fig. 3c**, **Fig. S9**). Asymmetric DMPE/PG also mimics the effects of a-DMPC/PG lipid mixtures, with addition of DMPE to the outer leaflet of DMPG liposomes accelerating folding and stabilising the inserted protein relative to symmetric DMPE:PG liposomes of equivalent outer lipid composition (**Fig 3b-c**). Interestingly, both DMPS and DMPE amplify the effect of asymmetry compared with DMPG and DMPC, perhaps due to the greater exposure of charges in these lipids. Asymmetric incorporation of DMPS and DMPE into liposomes thus has similar effects on OmpA as DMPG and DMPC, supporting the importance of lipid charge distribution in modulating folding. Excess negative charge in the outer leaflet slows folding while excess negative charge in the inner leaflet accelerates folding, presumably by raising and lowering the kinetic barrier to membrane insertion, respectively.

**Figure 3:**
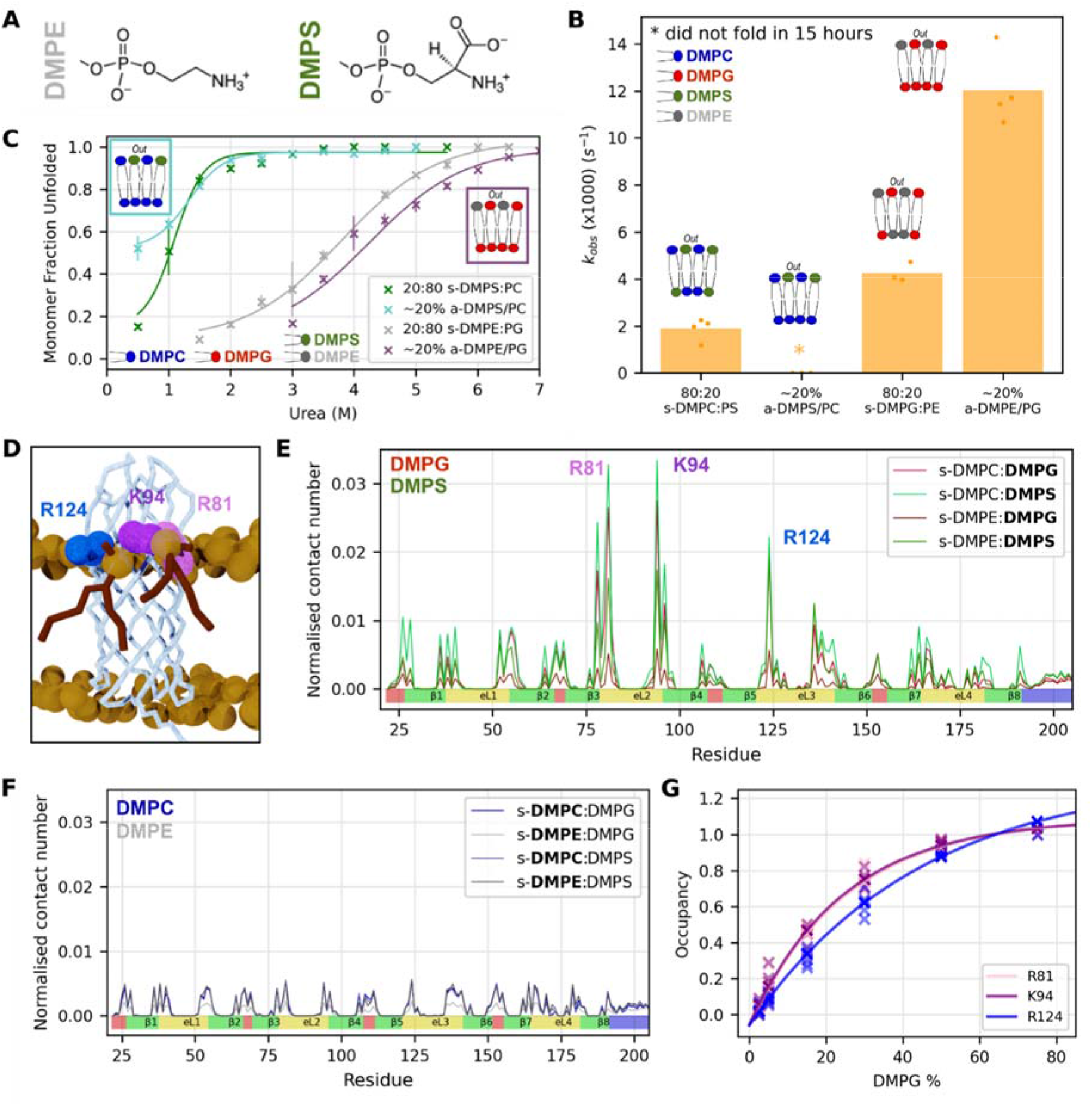
OmpA-lipid charge interactions dictate OmpA folding kinetics and efficiency. **(A)** Structure of DMPE (grey) and DMPS (green) headgroups (colours used throughout), charge analogues of DMPG and DMPC. **(B)** OmpA folding rate constants (s^-1^) into a-DMPS/PC or a-DMPE/PG LUVs and the equivalent symmetric liposomes (with the same outer leaflet content). * Indicates the folding had not finished in 15 hours (< 75% folded). **(C)** OmpA urea-titration folding curves into DMPC/PS and DMPE/PG symmetric and asymmetric liposomes. The data are fitted to the average ((line and crosses, 20% a-DMPS/PC line added to guide the eye but amplitude change too low to accurately fit).). **(D)** Final frame from an OmpA s- DMPG:DMPC simulation, showing two DMPG molecules (red) in the outer leaflet interacting with OmpA at R81, K94 and R124 (space fill in different colours). **(E)** Normalised contact count between OmpA and negatively charged lipids DMPG (red) or DMPS (green), and **(F)** between OmpA and zwitterionic lipids DMPC (blue) or DMPE (grey). Only the transmembrane region of OmpA is shown. Contacts are calculated over an average of 5 replicates. The secondary structure of the OmpA β-barrel is shown below (strands (green), extracellular loops (yellow), intracellular turns (red) and 14 residues of the periplasmic soluble domain (blue). **(G)** DMPG occupancy (fraction of time DMPG interacts with R81, K94 or R124) at different ratios of DMPC:DMPG. Data for 5 replicates are shown.

### The extracellular loops of OmpA specifically interact with negatively charged lipids

To further explore these effects, coarse-grained molecular dynamics (CG-MD) was used to search for interactions between folded (native) OmpA and lipid (**Table S1**), for a range of symmetric membranes of different DMPC-PG/PS ratios (**Fig. S10**-**11**). These simulations identified specific interactions between the head groups of DMPG and DMPS (but not DMPC or DMPG) and three positively charged residues (R81, K94 and R124) in the extracellular loops of OmpA (**Fig 3d-f**), with *in silico* mutation to serine removing these interactions (**Extended Data Fig. 7A**). The interaction time of DMPG with R81, K94 and R124 also depends on DMPG concentration (**Fig. 3g**) and suggests that a low binding site occupancy at lower DMPG concentrations may explain why OmpA folding is retarded at low DMPG concentrations (**Fig. 2a**). Simulations of BamA with s-DMPC:DMPG membranes also showed specific interactions between DMPG and residues K507, H533, K566, S764 and K793 in its extracellular loops (and K580 in an intracellular turn) (**Extended Data Fig. 7B**). These results support the view that charge-mediated lipid-protein interactions play an important role in the folding rate and apparent stability of OMPs.

### Lipid-OmpA charge interactions modulate folding

The extracellular loops of OmpA have seven positively charged Lys/Arg residues (R81, K85, K94, R124, K128, K134, R177) and seven negatively charged Asp/Glu residues (D41, E53, E89, D126, D137, D170, D179), most of which are highly conserved (**Extended Data Fig. 8**), including the three lipid-interacting positively charged residues (R81, K94 and R124) identified above. To explore the role of OMP-lipid charge interactions further, four variants of OmpA were created that differ in their extracellular loop charge: OmpA-NP (No Positive charges); OmpA-NN (No Negative charges; OmpA-NC (No Charges) (see Methods for these sequences); and OmpA-M3 (R81S, K94S & R124S). The folding rate and apparent stability of these variants folding into symmetric and asymmetric liposomes was then characterised (**Fig. 4a-d**, **Extended Data Fig. 9**).

**Figure 4:**
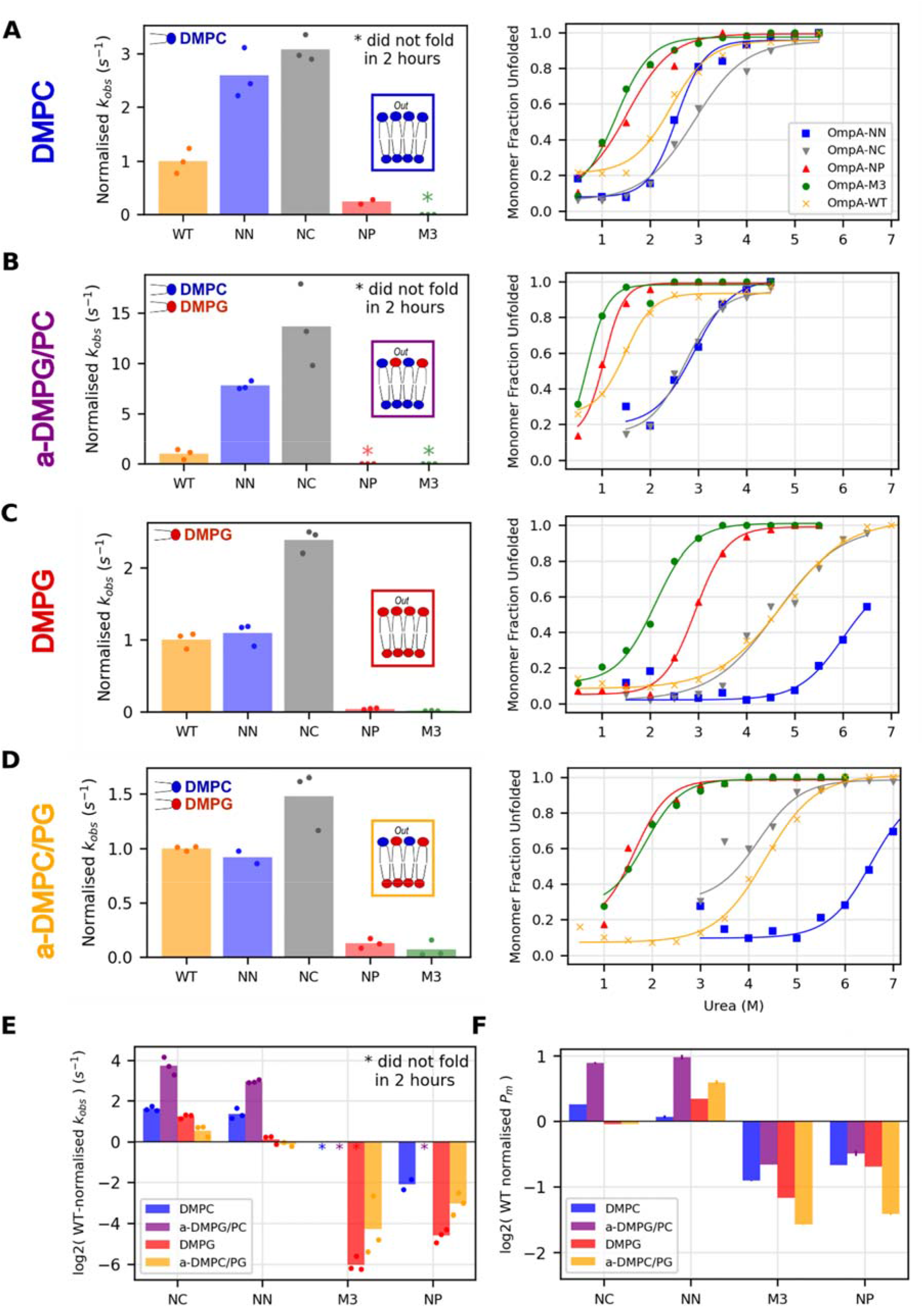
Folding kinetics and stability of OmpA charge variants compared to WT-OmpA. The relative folding rate constant difference (normalised to WT) and urea-titration folding curves for OmpA variants measured using cold SDS PAGE in **(A)** DMPC, **(B)** a-DMPG/PC PC (fit for OmpA-M3 urea folding added to guide the eye but amplitude too low to accurately fit), **(C)** DMPG or **(D)** a-DMPC/PG LUVs. OmpA-NC fraction folded at 3.5 M urea was excluded from the fit in (D). **(E)** Comparison of the average folding rate constants normalised to WT-OmpA (* indicates folding not complete (< 75% folded in 2 h). **(F)** P_ms_ (relative to that of WT OmpA) for all OmpA variants, error bars indicate +/- goodness of fit (reduced χ-squared).

The effects of the different lipid mixes on folding rate constant and stability of the different variants show similar trends (**Fig. 4e-f**). OmpA-NC folded the fastest in all liposome systems, likely because the uncharged extracellular loops can cross the bilayer most easily. OmpA-NN, which lacks negative charge, folds more rapidly than WT-OmpA in DMPC and a-DMPG/PC liposomes. In marked contrast, removing some, or all, of the positive charge (OmpA-NP and OmpA-M3) slowed folding at least 6-fold compared with OmpA-NC in the same lipid mixtures. For OmpA- M3, this effect is striking, ablating folding over the 2 h timescale of these experiments in two of the four liposomes tested (see asterisks in **Fig. 4a-f**) and in two instances having a greater effect than removing all charges in OmpA-NP. P_m_ values also follow this pattern, indicating that extracellular positively charged residues also enhance protein stability. Thus, regardless of lipid composition, the positively charged residues in the extracellular loops, especially at positions 81, 94 and 124 of OmpA, enhance the rate of folding and increase stability, and an asymmetry in charge across the bilayer modulates this effect, demonstrating an inter-dependence between protein and lipid charge in determining folding rate and stability.

### OMPs have a conserved patch of charged residues in their extracellular loops

The preference for positive charges in the extracellular loops of OMPs has been noted before ^39^, but to investigate this phenomenon in more detail, we examined the distribution of positively charged residues in the transmembrane OMPs from the Orientations of Proteins in Membranes (OPM) database, which were clustered by sequence to 75 proteins to reduce over-representation of close homologues (Methods). The spatial enrichment of different residues at each site was then calculated in 1 Å slabs parallel to the membrane plane, an analysis which identified the well characterised aromatic girdle ^39^ that flanks the hydrophobic acyl chains on each side of the membrane (**Fig. 5a**). However, minimal patterns of charged residues were seen in loop regions in this analysis, presumably because different residue probabilities in the transmembrane and water-soluble regions of the protein skew the statistics. To overcome this, the probability of residues being enriched in the extracellular loops was analysed against only the soluble regions of these OMPs (transmembrane residues excluded from the analysis) (**Fig. 5b**). This analysis revealed a patch of (>2σ significant) positive residues 6-10 Å above the plane of the membrane’s outer leaflet, matching perfectly the location of OmpA’s M3 group of residues (**Extended Data Fig. 12**).

**Figure 5:**
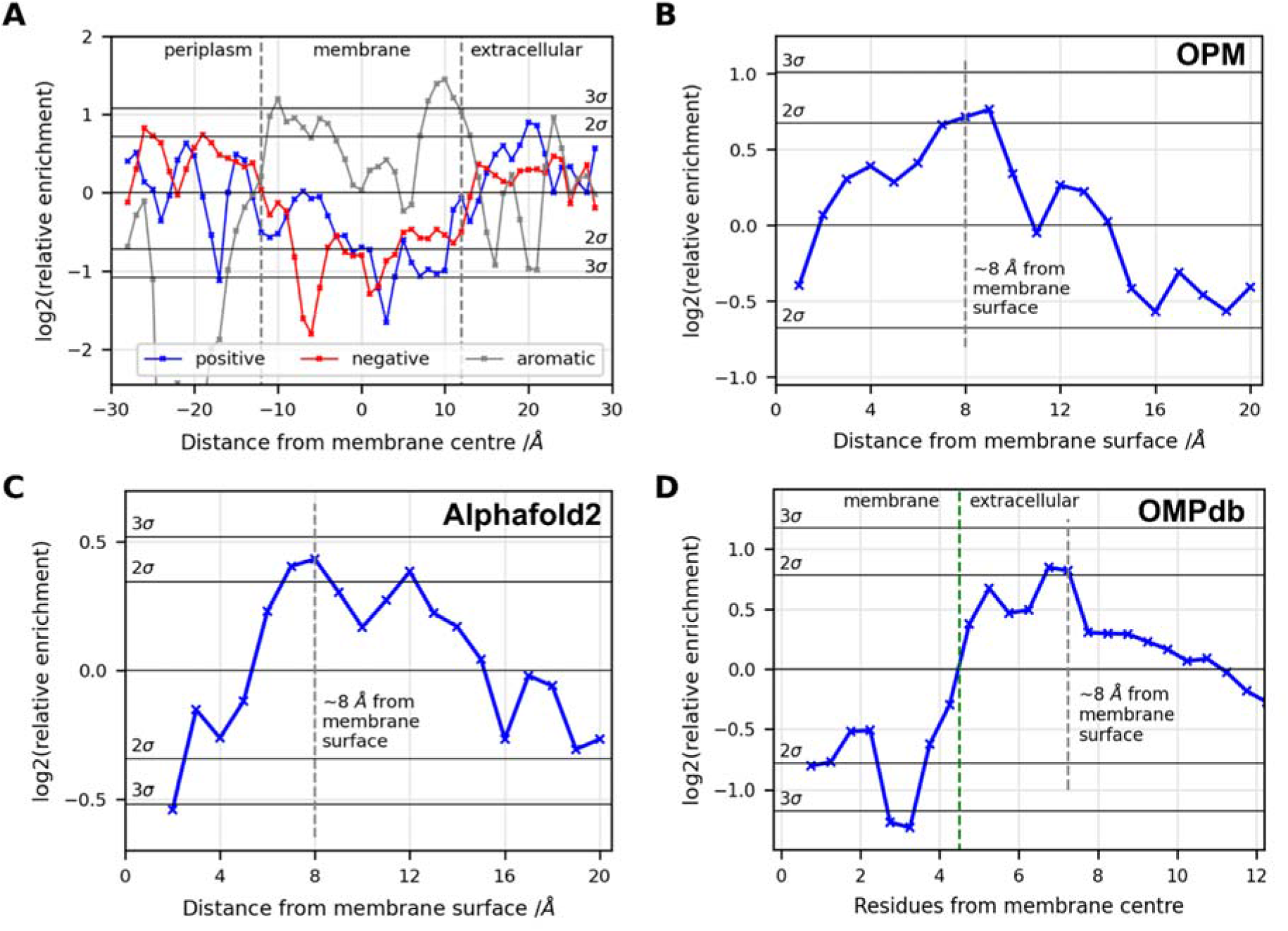
OMP residue enrichments perpendicular to the membrane plane show a conserved enrichment of positively charged residues in the extracellular loops ∼ 8 Å from the membrane surface. **(A)** Residue enrichments of aligned OMPs from the OPM database relative to the probability of finding an amino acid randomly, calculated over the whole protein sequence. Membrane thickness is the average of all OPM structures. **(B-D)** Residue enrichments in the extracellular loops of OMPs relative to finding an amino acid from the soluble regions of the protein randomly, calculated from **(B)** the OPM, **(C)** Alphafold2 and **(D)** OMPdb datasets.

While the OPM database is information rich, it contains relatively few OMPs. We therefore looked at OMP structures recently predicted by Alphafold2 ^62, 63^, and OMP sequences from the OMPdb^64^. A total of 2285 OM-annotated proteins were identified, and quality filtered based on signal peptide and transmembrane prediction, and Alphafold2 pLDDT score (Methods). Again, sequences were clustered to remove close homologues yielding 343 transmembrane OMPs. The OMPdb database was filtered by sequence identification (pHMM coverage) and topology prediction (Methods), and sequence clustered to 19,055 OMPs. These sequences do not contain explicit structural information, but topology predictions and an OPM-calibrated correlation curve between number of residues and distance from the membrane plane, allow residue count from the membrane to be used as a proxy for distance (**Fig. S12**). These analyses also show a peak (≥2σ) for enrichment for positive residues ∼8 Å from the membrane surface (**Fig. 5c-d**). No consistent pattern over all four datasets is observed for negatively charged residues (although a pattern of enrichment of negative charge residues was observed using Alpahfold2 and OMPdb) (**Fig. S12**). Collectively, these analyses identify the enrichment of positive residues in the extracellular loops ∼6-10 Å from the membrane surface. Given how strongly such positive charges impact the folding of OmpA, we suggest that these charges may be conserved to facilitate efficient OMP folding into the OM.

## Discussion

The folding kinetics of OMPs have been extensively studied, with more than 35 studies published on 15 different proteins over the preceding 30 years (reviewed in ^23, 26^). Despite this extensive body of literature, and the fact that an asymmetric distribution of lipids between the two leaflets of a bilayer is the norm for biological membranes, here we present the first systematic study of the folding of an OMP into asymmetric bilayers. The results are striking: asymmetric distribution of lipids between the two leaflets of a bilayer has a profound effect on both the observed rate constant for protein folding and on the apparent stability of the protein in the bilayer. This effect is mediated by the distribution of charge. Increasing the number of negatively-charged lipid head groups (either DMPG or DMPS) in the inner leaflet of the liposome (functionally equivalent to the outer leaflet of the OM), progressively reduces the kinetic barrier for folding and thus increases its rate. However, when negatively charged lipids are only present in a liposome’s outer leaflet (equivalent to the inner leaflet of the OM), stabilising lipid head group interactions with the natively folded protein in the inner leaflet cannot occur, with the result that OmpA (and BamA) do not fold (or do so only very slowly).

The results from liposomes composed of DMPE-DMPG are particularly interesting, as these are the dominant lipids in the inner leaflet of the bacterial OM ^65^. The symmetric incorporation of DMPE into DMPG liposomes slowed folding, consistent with studies in symmetric DMPC-PE mixes ^31^. However, we show this inhibition can be partially overcome by introducing the DMPE asymmetrically to the liposome outer leaflet only. Thus the reduction in folding rate is mediated by lipids in both leaflets, and is not simply a result of interactions at the liposome surface. Our results suggest that the asymmetric presence of the negatively charged LPS in the bacterial outer membrane could directly facilitate folding. Collectively therefore these data identify a key feature of OMP folding, namely that the balance of negative charge in both leaflets of a bilayer is critical for folding and could be modulated biochemically, forming a rheostat that tunes the folding rate by more than three orders of magnitude in the conditions sampled here. Notably, the lack of such charge asymmetry in the bacterial IM would retard aberrant folding into the IM and hence play a role in determining flux of OMPs to the OM.

Charge interactions within a membrane protein’s sequence play an established role in their folding ^35^, and the ‘positive outside’ rule, first described in 2005 ^38^, is a recognised feature of OMP sequence/structure. Here we reveal the molecular detail that underpins this phenomenon. Using MD simulation and mutational analysis we identify that a patch of external positive (PEP) residues within the external loops of OmpA, are critical for productive folding, rather than a general requirement for positive charge. These PEP residues lie ∼6-10 Å from the membrane surface and mediate OMP folding via interactions with the excess negative charge that we have identified above as being a key driver for efficient folding. We then show that this PEP is present in OMP sequences generally, suggesting that this feature may be a conserved determinant of efficient OMP folding. For the studies in liposomes this excess negative charge is the inner leaflet, but in the OM, the protein would approach the membrane from the periplasm, and excess negative charge particularly on LPS molecules, would be in the outer leaflet of the OM. Thus natural selection on OMP sequences and the machinery for the generation and maintenance of lipid asymmetry might plausibly operate synergistically to maximise the efficiency of OMP folding.

In summary, the results presented here not only provide new insights into how bilayer charge asymmetry effects the folding and stability of OMPs, but they also suggest roles of lipid asymmetry in a broader context in OMP folding. Specifically, by exploring the effect of lipid charge asymmetry with the model proteins OmpA and BamA we reveal charge-mediated features in both the lipid environment and protein sequences that reduce the kinetic barrier to folding and stabilise the final, membrane inserted state. Although the exact nature of the modulation will require more targeted studies of a broad range of OMPs, including lipid mixes incorporating LPS, these results provide unprecedented insights into how membrane charge asymmetry affects OMP folding and stability, suggesting new routes to manipulate OMP behaviour for biotechnology applications, and suggesting how cells might exploit lipid asymmetry to modulate the efficiency of OMP folding into the highly asymmetric bacterial OM.

## Acknowledgements

We thank all members in the Radford, Ranson & Kali groups for their helpful discussions and critical reading of this manuscript, especially members of the OMP group in Leeds. J.M. is funded by Wellcome (222373/Z/21/Z). We thank Nasir Khan for his excellent technical support, Jim Horne for support with molecular biology, Leon Willis for his help with DLS, Algy Kazlauciunas for help with measurement of the zeta-potential and Dheeraj Prakaash for help with MD. CryoEM imaging was carried out at the Astbury Biostructure Laboratory, which was funded by the University of Leeds and Wellcome (108466/Z/15/Z & 221524/Z/20/Z). SER holds a Royal Society Professorial Research Fellowship (RSRP\R1\211057). For the purpose of Open Access, the authors have applied a CC BY public copyright licence to any Author Accepted Manuscript version arising from this submission.

## Contributions

All authors designed the experiments. J.M. performed all of the research. All authors contributed to analysis of the data, and all authors wrote or edited the manuscript.

## Data Availability

Source data files are provided with this paper (all folding kinetics, EM images, gels/TLCs, DLS and ζ-potential data, thinned MD trajectories) and are freely available at the University of Leeds Data Repository: https://doi.org/10.5518/1168

## Materials and Methods

### Liposome preparation

DMPC, DMPG, DMPE and DMPS lipids (Avanti polar lipids) were prepared as stocks with concentrations of 25 mg/ml in chloroform. Liposome preparations were all made to ∼40 mM lipid concentration. Lipids were placed into amber glass vials and dried under N_2_, vacuum desiccated for >3 h and resuspended in buffer (10 mM Tris-HCl, 100 mM NaCl, pH 8.5). Following complete resuspension, samples were freeze-thawed five times in liquid N_2_ and a 42 ᵒC water bath, then extruded 31-times through 100 nm nucleopore polycarbonate track-etched membranes (Whatman; GE Healthcare) (Avanti extruder) at a temperature ∼10 ᵒC higher than the T_m_. DMPE lipids were sonicated rather than extruded. 1% (*mol/mol*) DPPE-rhodamine (Avanti) was introduced as a fluorescent label where indicated. Liposomes were used within 48 hours of their synthesis. Lipid concentrations of DMPC and DMPG liposomes were determined using absorbance, calibrated by the Stewart assay ^66^ (**Fig. S13**): samples were dissolved in 750 μl chloroform, to which 750 μl of guanidine ferric-thiocyanate were added (0.4 M guanidine thiocyanate, 0.1M iron (III) chloride hexahydrate). Samples were vortexed vigorously for 1 min. Following phase separation, the chloroform phase was removed with an 18-gauge needle, its absorbance at 448 nm measured and lipid concentration determined using the calibration prepared (**Fig. S13**).

## Lipid exchange

The following protocol was adapted from the previous publication (104). Concentrations of donor (Cd) and acceptor (Ca) lipid were determined by:

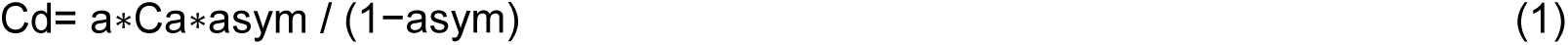

where *a* is the fraction of lipid accessible (∼0.5) and *asym* is the desired asymmetry (up to about 0.55). The concentration of MβCD (Cm) was determined by:

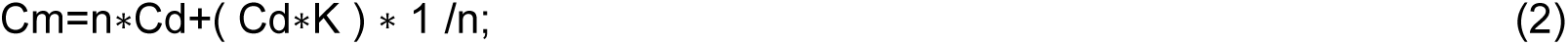

where n (set as 4) is the stoichiometry of the CD:lipid complex and K is an experimentally derived value (set as 292 M^-3^ for DMPC, DMPG and DMPS donation, and empirically adjusted to 150 M^-3^ for DMPE donation). These values are sensitive to MβCD activity and phospholipid-specific differences can be significantly reduced by using intermediate MβCD-lipid saturation (fixed at 60%). Donor liposomes were first solubilised with MβCD (Sigma) at 50 ᵒC, 1000 rpm for >20 min. Acceptor liposomes were then added and incubated at 35 ᵒC, 400 rpm for >20 min to allow for exchange. Liposomes were purified by two rounds of ultracentrifugation (105,000*g*, 4 ᵒC,30 min, Beckman Coulter, Optima MAX-XP). Following resuspension, liposomes were centrifuged at 5,000*g* for 5 min to remove aggregates. To ensure high sample yields only a single round of exchange was carried out, limiting asymmetry to ∼30% DMPC/PG and ∼ 50% DMPG/PC. Generated asymmetric liposomes were grouped to the nearest 10% (+/- 3%) for analysis. Exchanged liposomes were quality checked and used the same day they were made.

## Liposome absorbance analysis

Liposome absorbance was measured between 300-600 nm using quartz cuvettes. Absorbance traces were deconvoluted using a custom script, that found the liposome and fluorophore concentrations that minimised the following function

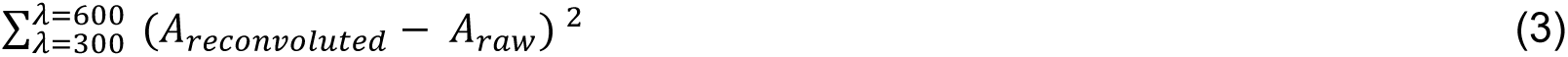

where A_raw_ is the raw absorbance trace and A_reconvoluted_ is the theoretical absorbance from the deconvoluted data.

## Determination of MβCD concentration using anthrone

Each sample (30 μl) was mixed with 100 μl anthrone reagent (0.2% (*w/w*) anthrone in 50% (*v/v*) H_2_SO_4_), heated at 95 ᵒC for exactly 10 min and then quenched by cooling on ice. Absorbance of the samples was measured at 630 nm. A calibration curve of 0-200 μM MβCD at 25 μM intervals was measured every time samples were assayed.

## Thin Layer Chromatography

Liposome samples were diluted to ∼0.5-2 mM and 5 µl dried under nitrogen. The sample was resuspended in chloroform and spotted on a TLC plates (Silica gel 60 F*₂₅₄*, Sigma 1.6834) and run with 40:9:6:3 (*v/v*) chloroform:methanol:ethanoic acid:water (DMPG-DMPC), 60:20:1 (*v/v*) chloroform:methanol:water (DMPG-DMPE) or 130:20:2 (*v/v*) chloroform:methanol:water (DMPC-DMPS). Plates were dried at 50 °C, dip stained into phosphomolybdic acid and developed by heating at 200 °C for exactly 20 min. Plates were imaged with a Q9 alliance imaging system (Uvitec) and densitometry analysed using ImageJ.

## ζ-potential and DLS

ζ-potentials and DLS were measured on a Zetasizer Nano ZS (Malvern) using DTS1070 cells at 25 ᵒC (60 sec incubation), 10-100 measurements were made in a water dispersant. Each sample was measured in triplicate, and cells were cleaned with 2% (*v/v*) Hellmanex, 18MΩ-H_2_O and then ethanol and dried under nitrogen. The cell quality was ensured using ∼5 measurements with a reference standard (DTS1235, Malvern).

## Imaging liposomes using cryo-EM

Samples (3 µl) of ∼0.5 mM liposomes were put onto glow discharged (PELCO easiGlow, Ted Pella Inc.) quantifoil grids (1.2/1.3) and incubated for 30 s. Grids were then blotted for 6 s with Whatman #1 filter paper at 4 °C and ∼ 90% relative humidity and plunge-frozen in liquid ethane using a Vitrobot Mark IV (Thermofisher). Grids were imaged on a 300 keV Titan Krios (ThermoFisher) electron microscope using EPU software and a K2 detector.

## ζ-potential prediction model

A review of the literature (89,104,126–137), combined with data presented here, yielded 315 data-points that met the following inclusion criteria: (i) T_m_ of all lipids of each sample must be known (except cholesterol which is handled separately; any liposomes without defined acyl chain composition are removed), (ii) measurement buffer salt must be NaCl or KCl, and (iii) ethanol must not be present in the buffer. Lipid composition of all the liposomes was parametrised with three values: (A) average overall charge per lipid; (B) average T_m_ of all lipids; and (C) fraction of lipid composition that is cholesterol.

An Extreme Gradient Boosted model (from XGboost library ^67^) was used with a root mean squared error loss function, a learning rate of 0.05 and an early-stop patience of 25 cycles (as evaluated from the current 25% validation data). The model was trained with the target ζ-potential using eight dataset features: salt concentration (monovalent), salt concentration (divalent), pH, hydrodynamic radii, temperature, overall charge, lipid T_m_, cholesterol fraction. The error associated with each measurement (the standard deviation value) was used to weight the features of an individual data-point, with weightings normalised between 0.375 and 0.625. Model hyper-parameters were explicitly optimised to reduce model over-fitting identified in early testing: subsample-per-node: 0.85, subsample-per-tree: 0.85, minimum-child-weight: 2.5 and maximum tree depth: 6. The models were validated with 4-fold cross-validation. Predictions were made by training a 50 model ensemble (all with mean average error (MAE) < 5 mV) on-the-fly and averaging their predictions to obtain a final value. The weight/gain-per-feature was analysed using python package scikit-learn. (Code can be accessed at: https://github.com/JonMarks29/zeta-potential-prediction)

## Plasmids and Mutants

Sequence alignment of OmpA homologs identified residue substitutions of charged residues within its extracellular loops. The most common alternative residue was used for generating the OmpA variants, or for residues that are completely conserved, they were substituted with serine. OmpA-NP: R81S, K85T, K94S, R124S, K128G, K134S, R177S; OmpA- NN: D41S, E53N, E89V, D126S, D137S, D180S, D189S; OmpA-NC: combination of both OmpA-NP and OmpA-NN; OmpA-M3: R81S, K94S, R124S. Genes encoding mutants of OmpA were ordered from GeneWizz, ligated into a pET11a vector using flanking BamHI and NdeI restriction sites and validated by sequencing.

## Protein expression and purification

Competent BL21(DE3) *E. coli* cells were transformed with the relevant plasmid (carbenicillin resistant), grown overnight at 37 °C on agar plates, a single colony picked and grown overnight in ∼20 ml of LB containing 100 µg/ml carbenicillin (37 °C, 200 rpm). 5 ml of culture was added to 500 mL of LB, grown to an OD_600_ of ∼0.6 and protein expression was then induced with 1 mM IPTG. Three hours post-induction, cells were harvested (5000 *g*, 15 min, 4°C) and the cell pellet frozen. After thawing the pellet was resuspended in 20 ml buffer (50 mM Tris-HCl pH 8.0, 5 mM EDTA, 1 mM PMSF, 2 mM benzamide), and the cells lysed via sonication. Following centrifugation (25,000 *g*, 30 min, 4 °C) the pellet was resuspended in 20 ml buffer (50 mM Tris-HCl pH 8.0 2% (*v/v*) Triton-X-100) and incubated for 1 h (room temperature, 50 rpm). Following centrifugation (25,000 *g*, 30 min, 4 °C) the supernatant and cell debris were removed from the resulting inclusion body pellet. The inclusion bodies were washed twice by resuspending in 50 mM Tris-HCl pH 8.0 and incubating for 1 h (room temperature, 50 rpm) before pelleting by centrifugation (25,000 *g*, 30 min, 4 °C). Inclusion bodies were solubilised in 25 mM Tris-HCl and 6 M Gdn-HCl (pH 8.0) for 1 h (60 rpm stirring), and following a final centrifugation (25,000 *g*, 30 min, 4 °C), the supernatant was loaded onto a Superdex 75 HiLoad 26/60 SEC column (GE Healthcare), equilibrated in 25 mM Tris-HCl pH 8.0, 6 M Gdn-HCl. Protein fractions were collected and concentrated to ∼100 μM (Vivaspin concentrators) and flash frozen for -80 ᵒC storage. Prior to folding, proteins were buffer exchanged into Tris-buffered saline (TBS, 20 mM Tris-HCl, 100 mM NaCl, pH 8.0), 8 M urea using Zeba™ Spin Desalting Columns, 7k MWCO, 0.5 ml (Thermo Scientific).

## Protein gel-electrophoresis

Samples were mixed 1:3 with loading dye (50 mM Tris-HCl pH 6.8, 6% (*w/v*) SDS, 0.3% (*w/v*) bromophenol blue, 40% (*v/v*) glycerol), boiled if required (>10 min, 100 ᵒC) and ∼14 μl loaded onto the gel. The molecular weight markers used were Precision Plus ProteinTM Dual Xtra Standards (BioRad). 15% Tris-tricine gels were homemade and contained 0.1 % (*w/v*) SDS, 1 M Tris-HCl at pH 8.45 and 13.3% (*v/v*) glycerol was included in the resolving layer. The cathode buffer was 100 mM Tris-HCl, 100 mM tricine, 0.1% (*w/v*) SDS, pH 8.25 and the anode buffer, 200 mM Tris-HCl, pH 8.9. A constant current of 30 mA (stacking) and 60 mA (resolving) was used. Following staining (InstantBlue® Coomassie, Abcam) gels were imaged with a Q9 alliance imaging system (Uvitec) and densitometry analysed using ImageJ. Fraction folded was calculated using only the monomer bands as folded/(folded+unfolded). (Inclusion of the higher order bands as unfolded species or by normalising folding against the boiled sample made no significant difference to the fraction folded, compared to ^68^, possibly due to the use of full length OmpA here). All WT-OmpA liposome conditions were tested in at least duplicate, all OmpA mutants were measured once.

## Determination of the intrinsic folding rates

The kinetics of intrinsic folding were measured using a QuantaMaster Fluorimeter (Photon Technology International (PTI)) controlled by FelixGX software v4.3, including a peltier-controlled temperature unit. Excitation/emission wavelengths of 280/335 nm were used. OmpA was buffer exchanged from 25 mM Tris-HCl and 6 M Gdn-HCl (pH 8.0) into 10 mM Tris-HCl and 8 M urea (pH 7.4) (Zeba spin desalting columns (Thermo Scientific)). Folding was initiated by rapid dilution of a 3.3 μM unfolded OmpA stock in 8 M urea to a final concentration of 0.2 μM OmpA and 0.48 M urea in the presence of 0.32 mM liposomes (LPR of 1600:1 (*mol/mol*) in 10 mM Tris-HCl and 100 mM NaCl at 30 °C. A minimum of three samples were measured for each liposome preparation, and the kinetics fitted to one-phase exponentials with a custom python script using SciPy to derive the observed rate constants which were then used for further analysis. Kinetics for DMPS containing liposomes were fitted to a two-phase exponential model based on high residual error in one-phase fits. Kinetic traces which folded to an amplitude of < ∼75% were not fitted.

## Measurement of protein stability by urea titration

Tryptophan fluorescence emission spectra (300-400 nm) with excitation at 280 nm were measured on samples that had been incubated overnight in different concentrations of urea at 30 °C to ensure equilibrium was reached. The fraction folded protein was then determined by taking the 335/350 nm ratio.

## Coarse-grained molecular dynamics (CG-MD) simulations

A structural model of full-length OmpA was predicted using Alphafold2, and for BamA the crystal structure (PDB 5D0O) was used. Following any *in silico* mutations, structures were coarse-grained using the martinize script. CGMD was conducted using gromacs (v5.0.7) ^69^ with Martini (v2) force field ^70, 71^. Bilayers were built around the protein using the insane script ^72^. CG waters were added and then the system neutralised with and 0.1 M of NaCl added. The system was energy minimised (steepest descent algorithm) and equilibrated with the protein backbone particles position-restrained for 3 ns. The equilibrated system was used to generate production systems for 3 μs (**Supp. Table 1**), with a 20 fs time-step and frames generated at 200 ps intervals. The barostat and thermostat were Parinello-Rahman (1 bar) ^73^ and V-rescale respectively ^74^. A compressibility of 3×10^-4^ bar^-1^ was used. The LINCS algorithm constrained bond lengths ^75^. Lipid protein contact analysis used a 0.55 nm distance cutoff to define contacts, performed on merged data from all replicas using gmx mindist. All lipid-protein contacts were normalised to lipid concentrations. For lipid density analysis the trajectories of all simulation replicas were concatenated and the protein orientation centered and fixed (gmx trjconv), gmx densmap was used to calculate densities. Residence time was calculated using pyLIPID ^76^.

## Laurdan assay

Lipid transition temperatures were measurement by laurdan fluorescence using a method adapted from ^61^. Laurdan, dissolved in DMSO, was added to preformed liposomes at a ratio 3200:1 *mol/mol* (lipid:laurdan) and a final DMSO concentration of 0.1 % (*v/v*). Liposomes were incubated near their transition temperature overnight. Laurdan fluorescence was excited at 340 nm and its emission at 440 and 490 nm measured for 10 s using a PTI fluorimeter as described above. Spectra were acquired at either 1 °C or 0.25 °C intervals at temperatures spanning approximately +/- 10 °C around the transition temperature, with 3 min equilibration at each temperature. General Polarisation (GP) was determined from the intensity at 440 nm and 490 nm (I, averaged over 10 s acquisition) using the equation

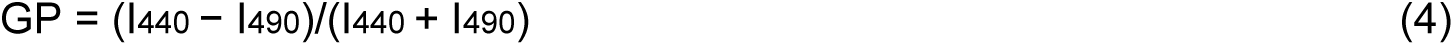

Midpoints were determined by numerically taking the first differential of the data.

## Bioinformatics

394 OM-annotated proteins from the Orientations of Proteins in the Membrane (OPM) ^77^, of which 198 have transmembrane regions, were sequence clustered to 70% sequence identity using CD-Hit ^78^ and manually inspected. Proteins are aligned in the membrane by OPM, 3D- space was split into 1 Å slabs parallel to the membrane plane, and residues assigned to slabs based on their Cα atom position, the number of residues per slab are indicated shown in **Fig. S14**. The enrichment/depletion of residues was calculated relative to the total amino acid content in the protein or in the soluble regions. 2σ/3σ significance was calculated separately for enrichment and depletion by finding the standard deviation of all positive and negative enrichments. See **Table S2** for a list of the proteins used.

2285 OM-annotated proteins were identified in the EBI-Alphafold2 database ^63^, (as of December 2021). Signal peptides were predicted using SignalP v5.0 ^79^ and removed from the structures (proteins with >90% prediction confidence taken forwards). Proteins were also filtered by Alphafold2’s pLDDT (>80%) leaving 1765 proteins. Transmembrane regions and membrane orientation was predicted using Immers ^77^ and 842 proteins identified with >0 transmembrane regions (693 >8 strands, i.e. are full barrels). Sequences were clustered to 70% sequence identity using CD-Hit ^78^, leaving 343 structures, which were processed as for the OPM dataset. See **Table S2** for a list of the proteins used.

The ∼1.3 x10^6^ sequences in the OMPdb (as of August 2021) ^64^ were quality filtered based on by topology prediction and pHMM coverage score (both >95%) and sequences missing residues removed, leaving 71, 181 sequences. These were sequence clustered to 70% sequence identity using CD-Hit ^78^, leaving 17, 931 sequences. Residue enrichment was carried out as above using residue count away from the centre of the membrane to split the protein into slabs. A distance calibration for residue count was determined from the OPM structures databases (**Fig. S12**). See **Table S2** for a list of the proteins used.

---

**Extended Data Figure 1:**
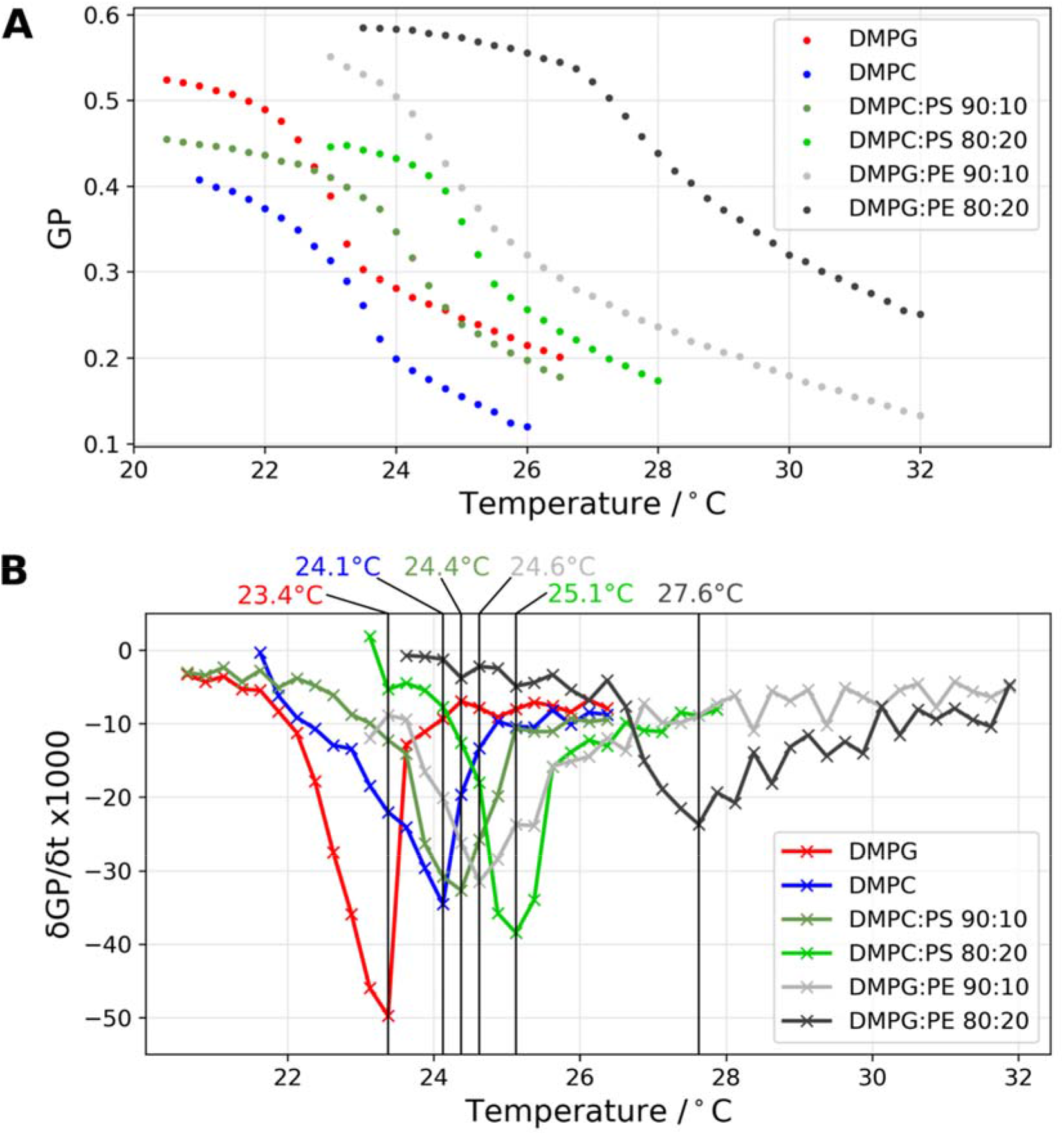
Global lipid phase transition behaviour for liposomes used in this study, measured using laurdan fluorescence. **(A)** The GP (generalised polarisation ratio of fluorescence at 440 and 490 nm, see Methods) against temperature for pure DMPC and pure DMPG liposomes, and DMPS-DMPC and DMPE-DMPG lipid mixes, as indicated, measured using 0.25 °C intervals. (B) The first derivative of the GP, with the implied Tms.

**Extended Data Figure 2:**
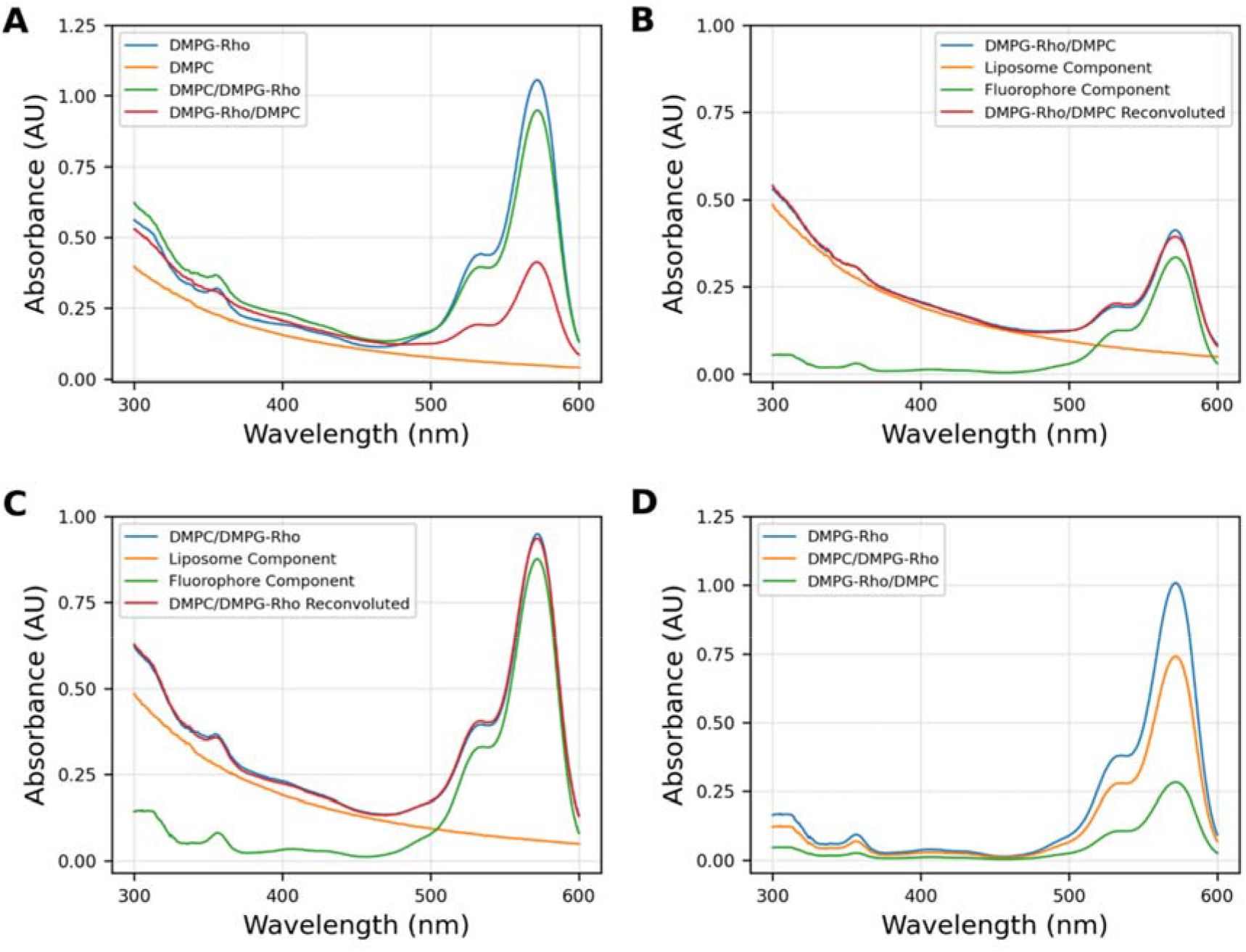
DMPC and DMPG lipids are competent to exchange through MβCD-mediated exchange. **(A)** Raw absorbance spectra of unexchanged DMPG-Rho liposomes (DMPG + 1% (mol/mol) DPPE-rhodamine) and DMPC liposomes, and exchanged liposomes, as indicated. **(B)** Deconvoluted absorbance spectra for DMPG-Rho/DMPC exchange and **(C)** DMPC**/**DMPG-Rho exchange, separating the liposome and fluorophore absorbance components. **(D)** Liposome concentration normalised fluorophore components of DMPG-Rho and the exchanged samples, showing loss of fluorescence from the DMPC/DMPG- Rho and gain of fluorescence in the DMPG-Rho/DMPC samples, indicating successful lipid exchange.

**Extended Data Figure 3:**
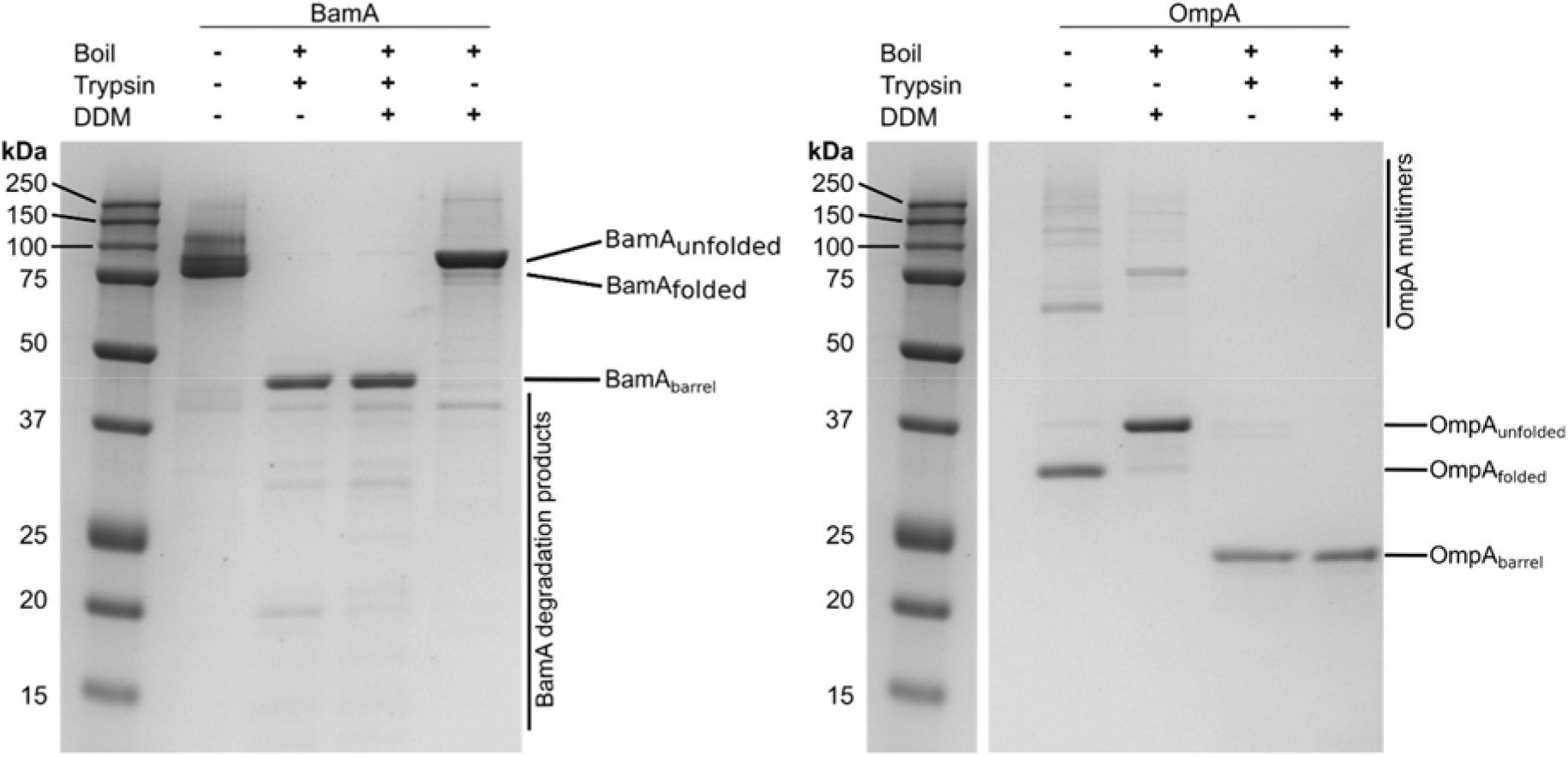
OmpA and BamA fold unidirectionally. OmpA and BamA were folded into DMPC liposomes and each were then incubated with trypsin (1:10000 molar ratio substrate:trypsin) overnight and compared to DDM-solubilised and trypsin cleaved samples treated identically. OmpA and BamA each show complete cleavage of their periplasmic domains, indicating that they have folded unidirectionally, with the water-soluble domains exposed to the bulk solvent.

**Extended Data Figure 4:**
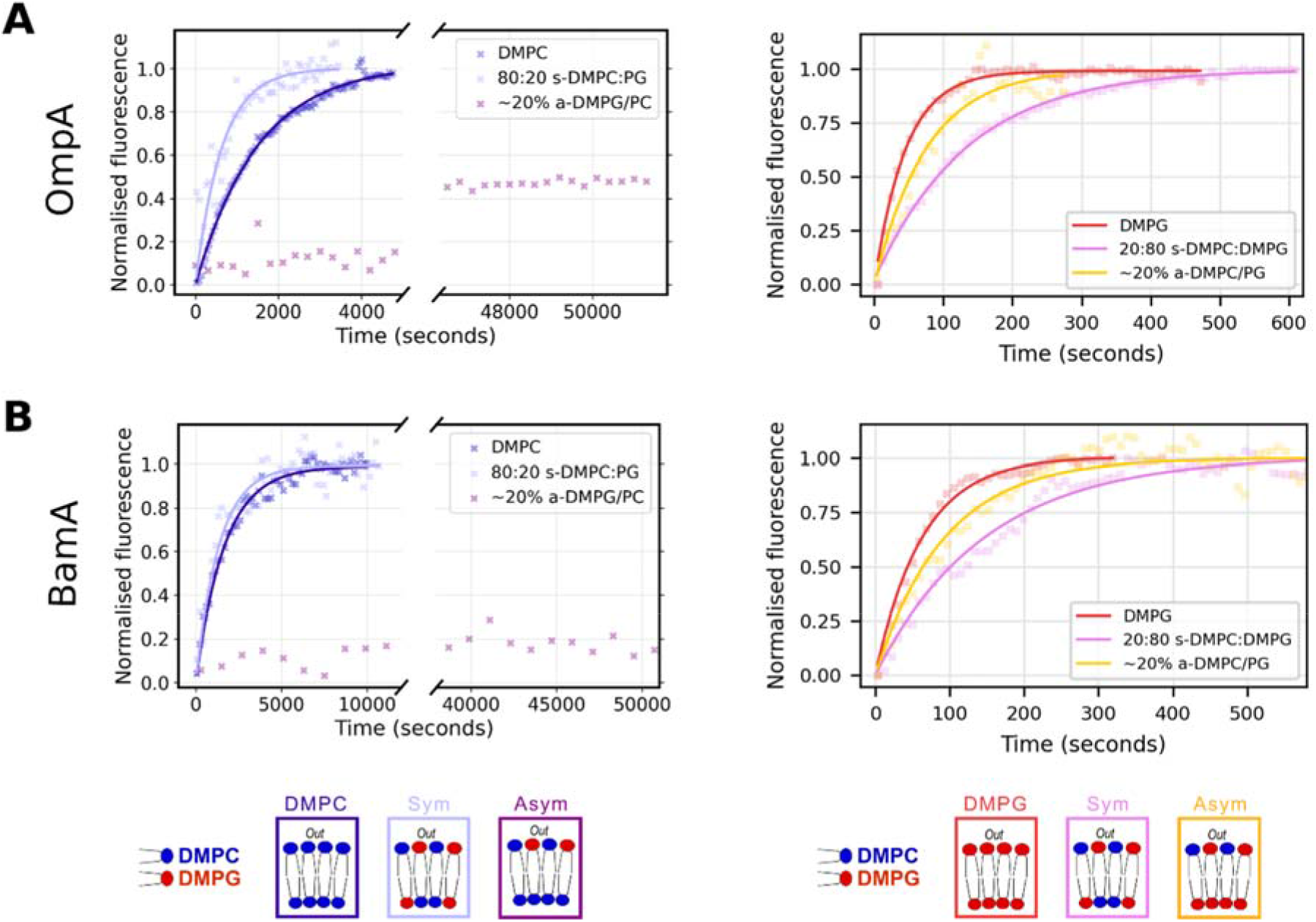
Example kinetic data and fits with representative lipid environments for folding of (A) OmpA and (B) BamA. Kinetic data shown for DMPC, DMPG and 20% symmetric and asymmetric liposomes. Data were normalised for comparison, ∼20% a-DMPG/PC did not reach completion for either OmpA or BamA and these data were normalised to their respective DMPC traces.

**Extended Data Figure 5:**
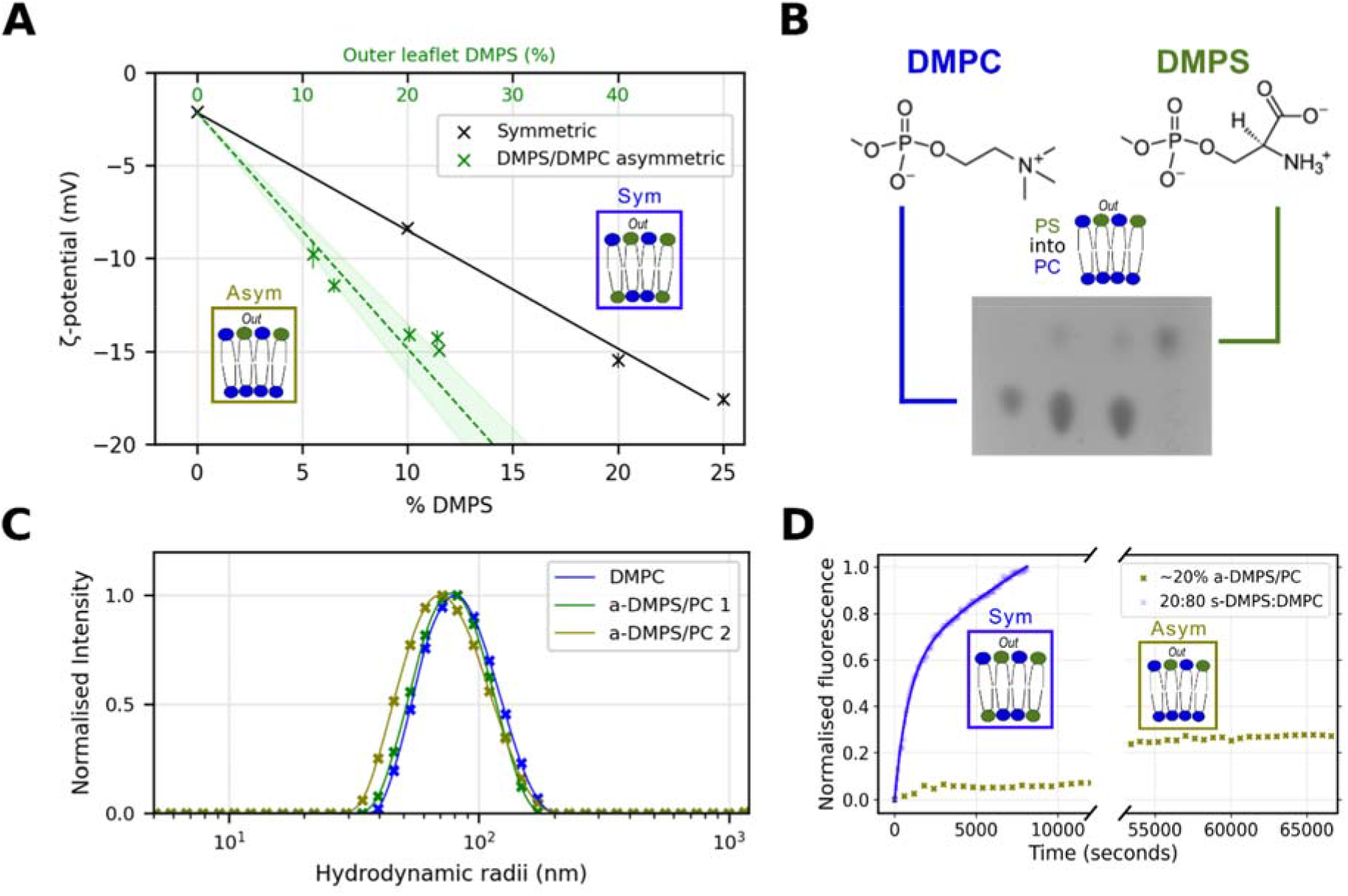
Generating asymmetric DMPS/DMPC liposomes. **(A)** ζ-potential calibration curve for asymmetric DMPS-DMPG lipid mixes, for DMPS fractions 0-25%, showing asymmetric liposomes (green crosses). **(B)** TLC of duplicate ∼20% a-DMPS/DMPC exchanged liposomes. **(C)** DLS of pre-exchange DMPC and duplicate post-exchange DMPS/DMPC liposomes. **(D)** Example OmpA folding kinetic traces, measured by tryptophan fluorescence, for 20:80 s-DMPS:DMPC (kinetic fit shown, blue line) and ∼20% a-DMPS/DMPC (did not complete folding (< 30% folded) over >15 hours, not fitted). Two-phase exponentials were fitted to DMPS containing liposomes.

**Extended Data Figure 6:**
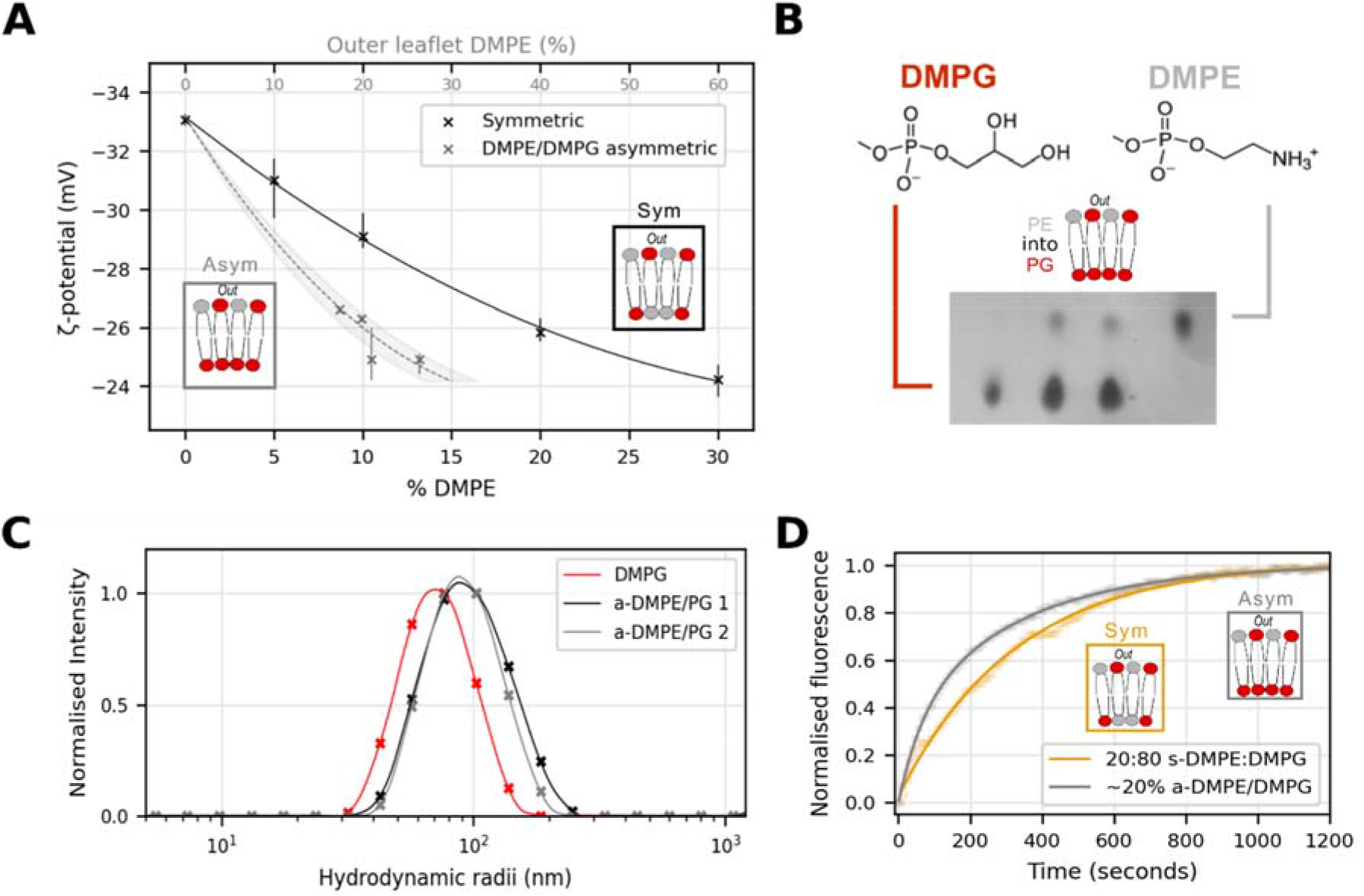
Generating asymmetric DMPE/DMPG liposomes. **(A)** ζ-potential calibration curve for asymmetric DMPE-DMPG lipid mixes, for DMPE fractions 0-30%, showing asymmetric liposomes (grey crosses). **(B)** TLC of duplicated ∼20% a-DMPE/DMPG exchanged liposome. **(C)** DLS of pre-exchange DMPG and duplicate post-exchange a-DMPE/DMPG liposomes, the slight increase in liposome size post-exchange is likely due to the readiness of DMPE to exchange into the liposomes. **(D)** Example OmpA folding kinetic traces, measured by tryptophan fluorescence, for 20:80 s-DMPE:DMPG (kinetic fit shown, yellow line) and ∼20% a- DMPE/DMPG (kinetic fit shown, grey line).

**Extended Data Figure 7:**
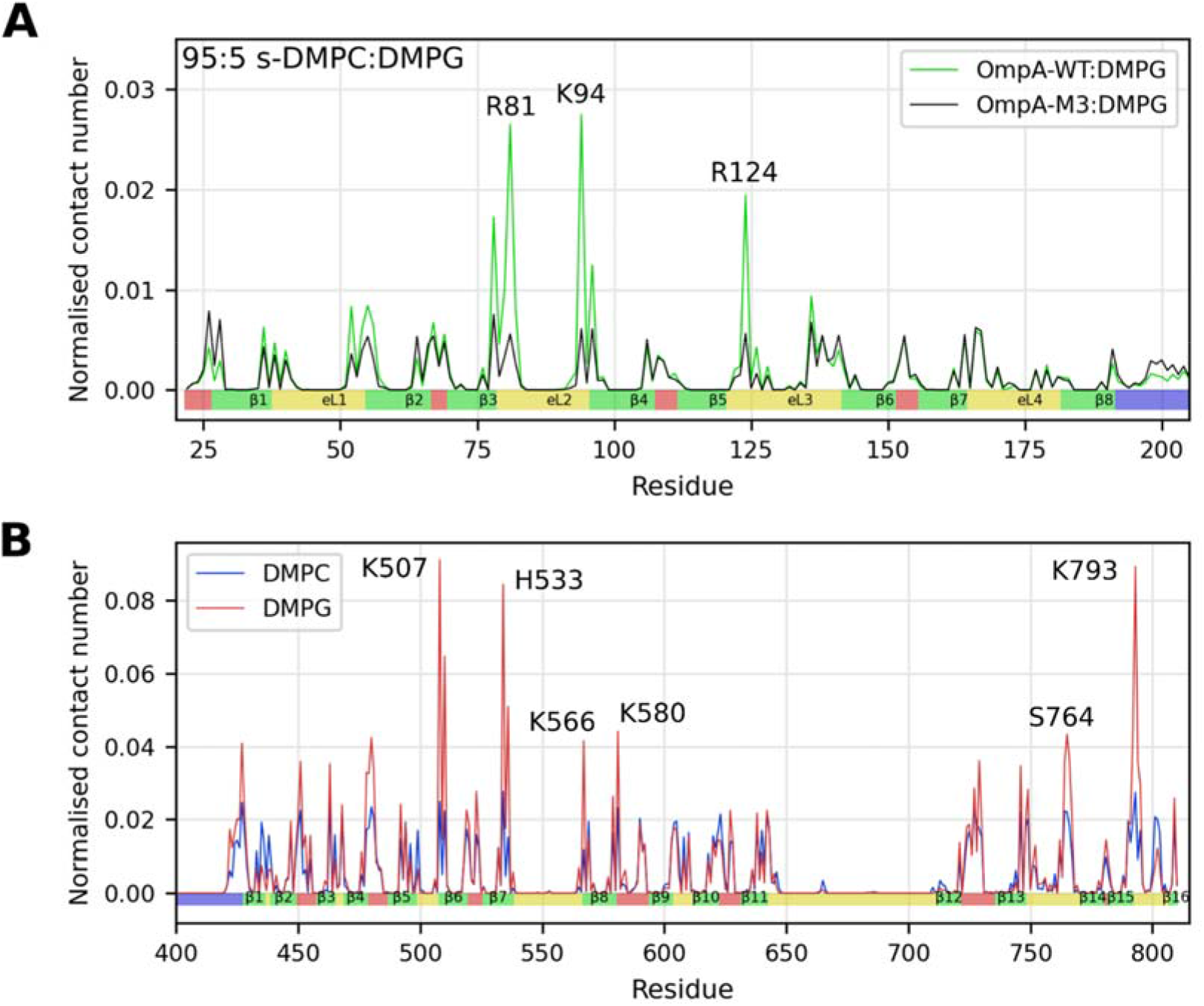
Normalised contact plots of *in silico* mutated OmpA-M3 (R81S, K94S and R124S) and WT-BamA to DMPG and DMPC lipids. **(A)** Normalised contact number for the transmembrane region of OmpA-WT and OmpA-M3 in a 95:5 s-DMPC:DMPG membrane. Contacts normalised by lipid concentration (Methods). Substitution of these three positive residues with Ser eliminates specific DMPG binding. (**B**) Full length BamA was simulated in a 95:5 s-DMPC:DMPG system. Only the transmembrane region is shown for clarity, structural features are shown at base of plot (strands (green), extracellular loops (yellow) intracellular turns (red) and 24 residues from POTRA5 (blue). Interactions with a normalised contact number >3σ are labelled, and indicated in main text.

**Extended Data Figure 8:**
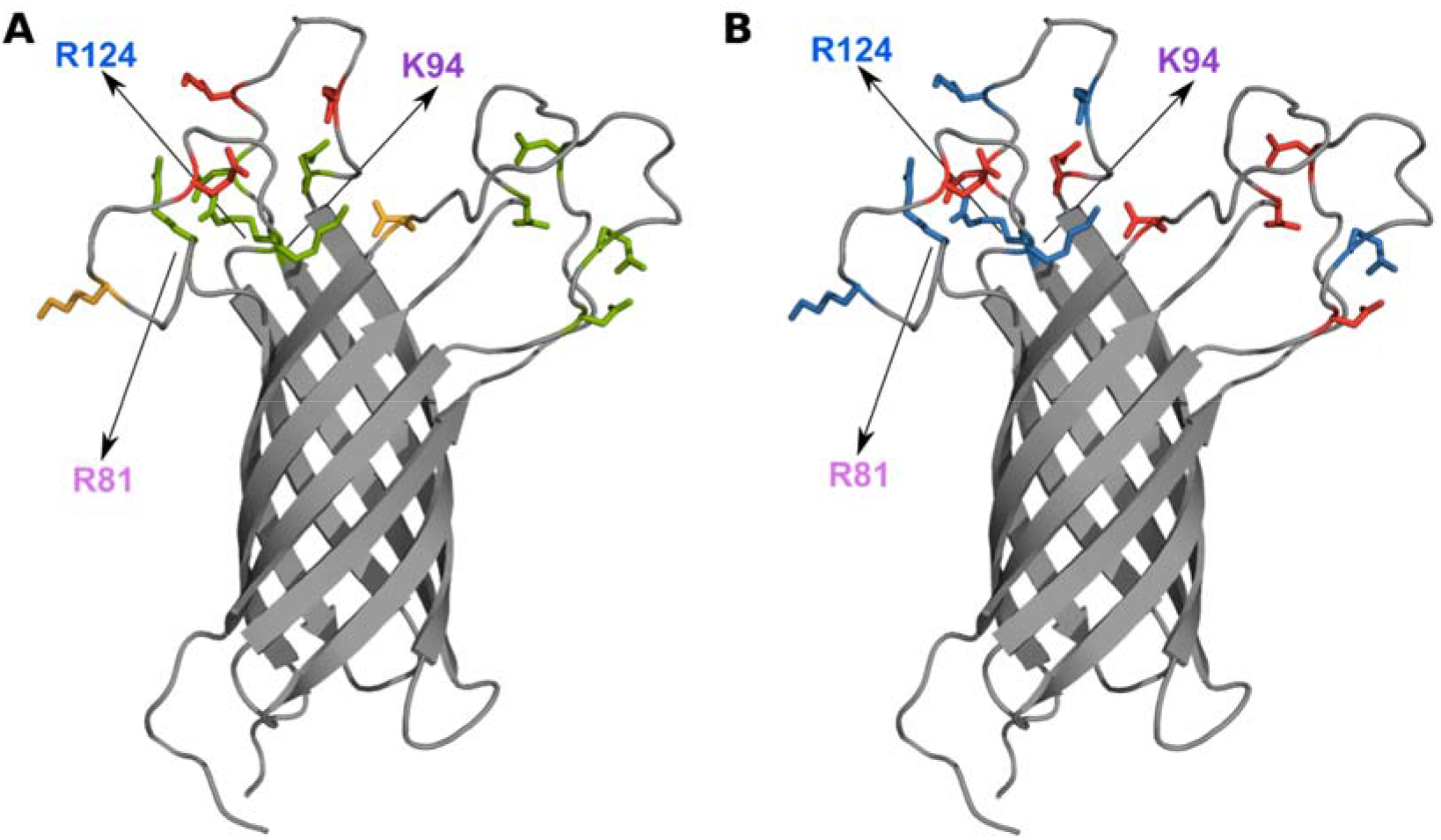
Conservation and location of Lys/Arg positively and Asp/Glu negatively charged residues in the extracellular loops of OmpA. **(A)** Relative conservation of charged residues (green: well conserved (>99%), yellow: partially conserved (>90%), red: less conserved (<90%)) over 2750 OmpA barrel sequence homologs. Conservation is the retention of a K/R or D/E at a given position. (**B**) Spatial distribution of positively (Lys/Arg) and negatively charged (Glu/Asp) residues in the extracellular loops of OmpA (blue: positive, red: negative) (PDB: IG90). R81, K94 and R124 that specifically interact with negatively charged lipid are labelled.

**Extended Data Figure 9:**
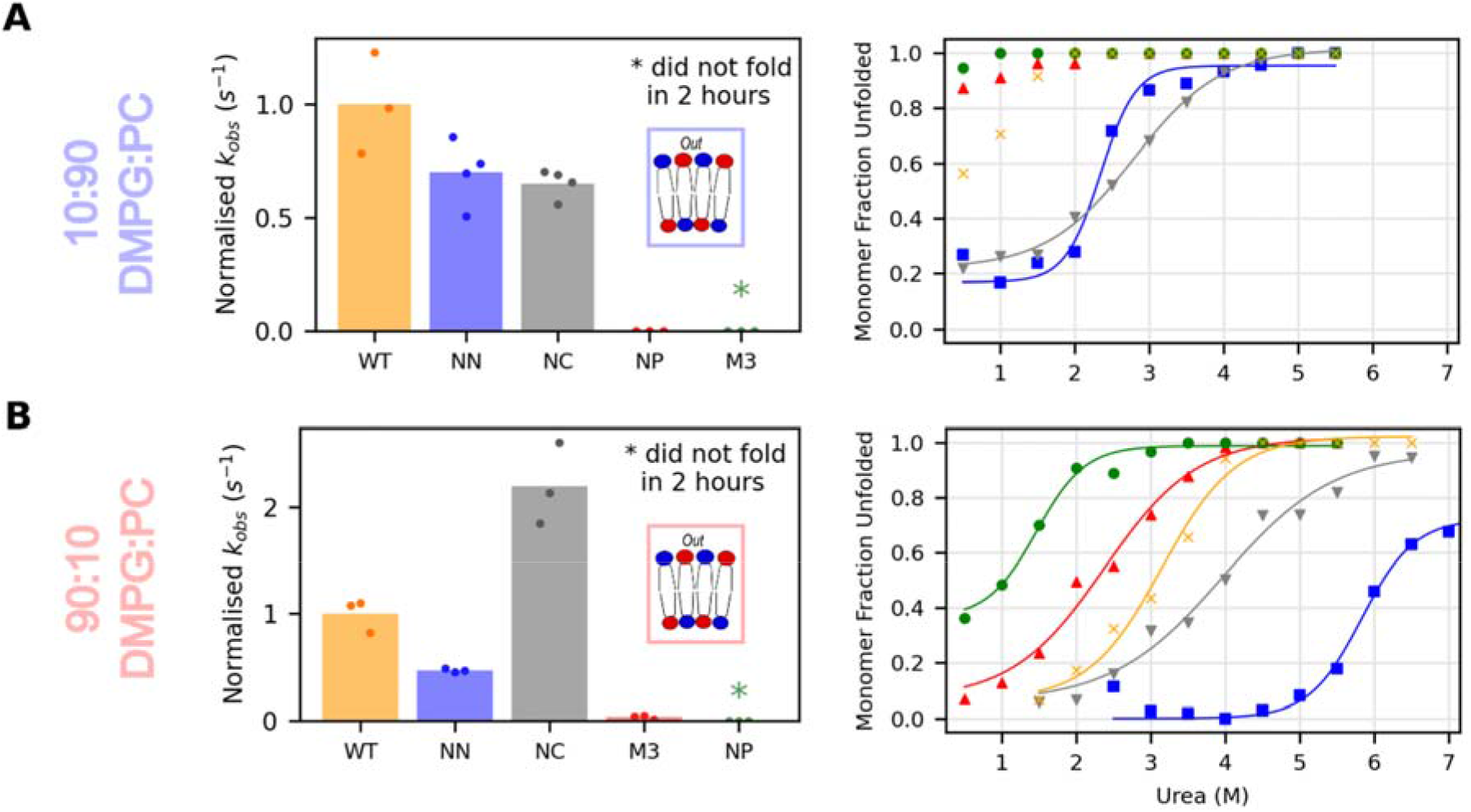
Folding kinetics and urea-titration of OmpA charge variants compared with WT OmpA in symmetric 90:10 and 10:90 DMPC:DMPG LUVs. The difference in folding rate constants (normalised to WT OmpA) and apparent stability vs. urea concentration for OmpA-variants in (A) 10:90 DMPG:PC and (B) 90:10 DMPG:PC symmetric LUVs are shown, as indicated in the key.

**Extended Data Figure 10:**
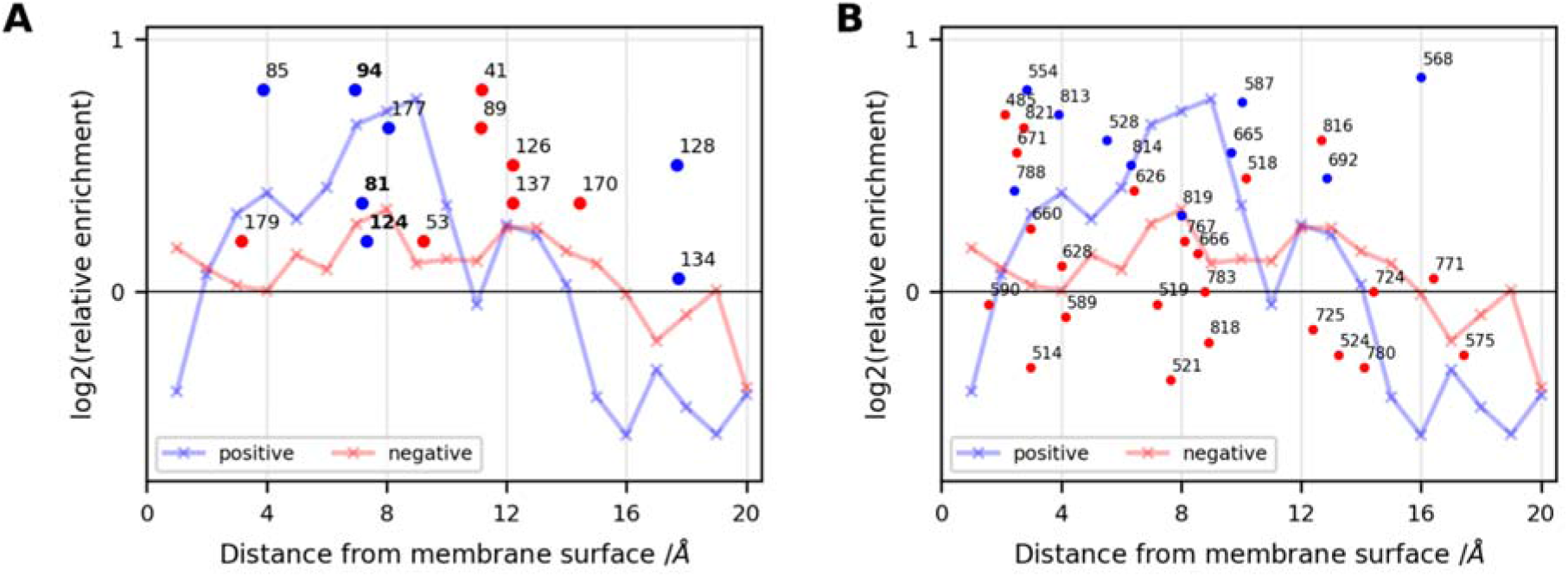
The extracellular charged residues of (A) OmpA and (B) BamA generally agree with the patterns of positive and negative residues identified by analysis of 199 sequences in the OPM (solid lines). Residues R81, K94 and R124 in OmpA are identified as lipid interacting by CG-MD (OmpA-M3 cluster), shown in bold. Positive residues are shown as blue circles, negative residues as red circles and are labelled with residue number above-right of the marker.

**Supplementary Figure 1:**
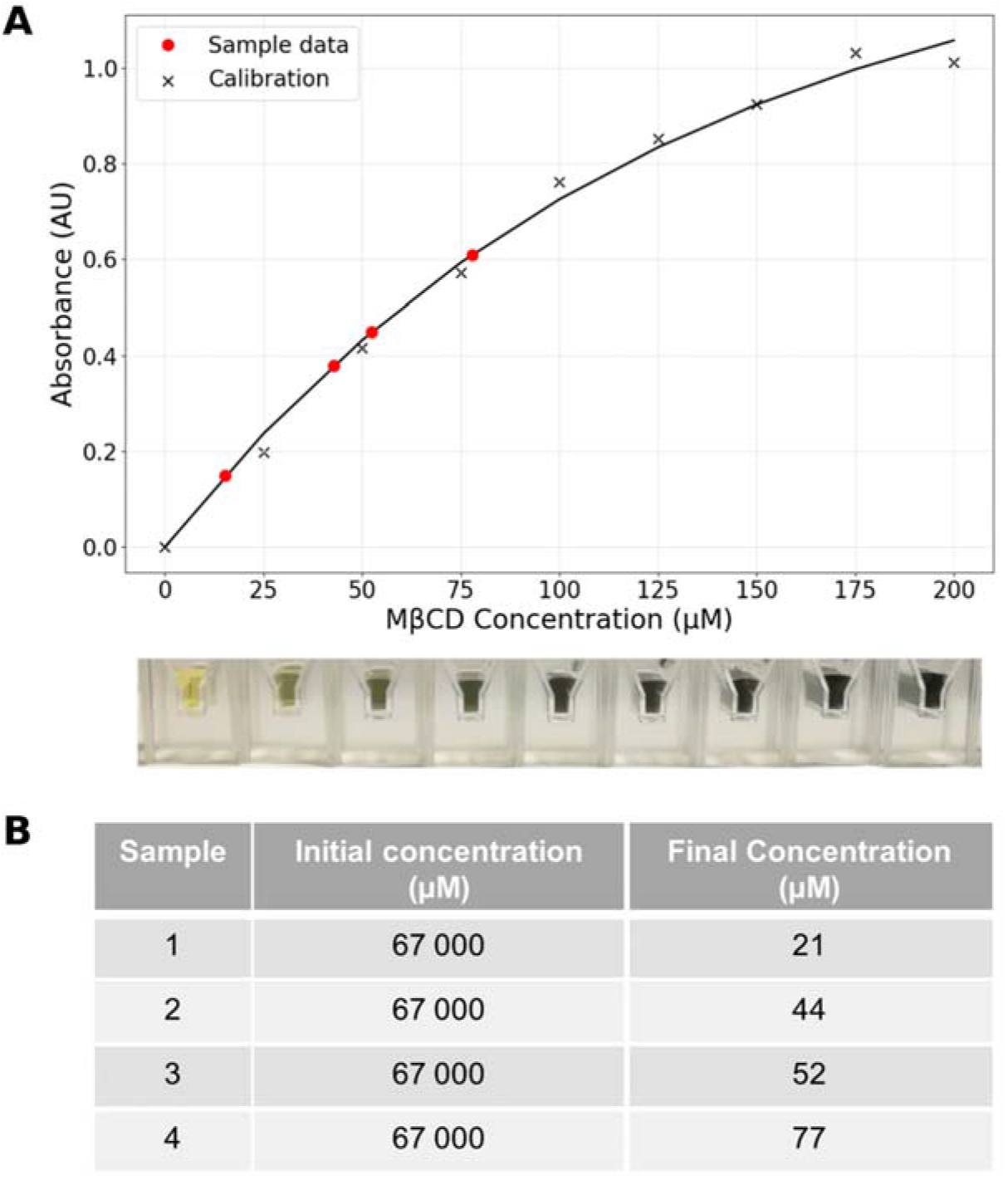
MβCD is effectively removed by ultracentrifugation. **(A)** The sugar anthrone assay can colorimetrically detect MβCD at µM concentrations (black crosses: calibration, red circles: sample data). Calibrant samples are shown below**. (B)** Example measurements of MβCD concentrations before and after ultracentrifugation (red circles in panel (A)). Typical reductions are >1000x, reducing free MβCD, and hence its associated lipid, to insignificant quantities for the assays described. Four replicates with an identical initial concentration of 67 mM are shown.

**Supplementary Figure 2:**
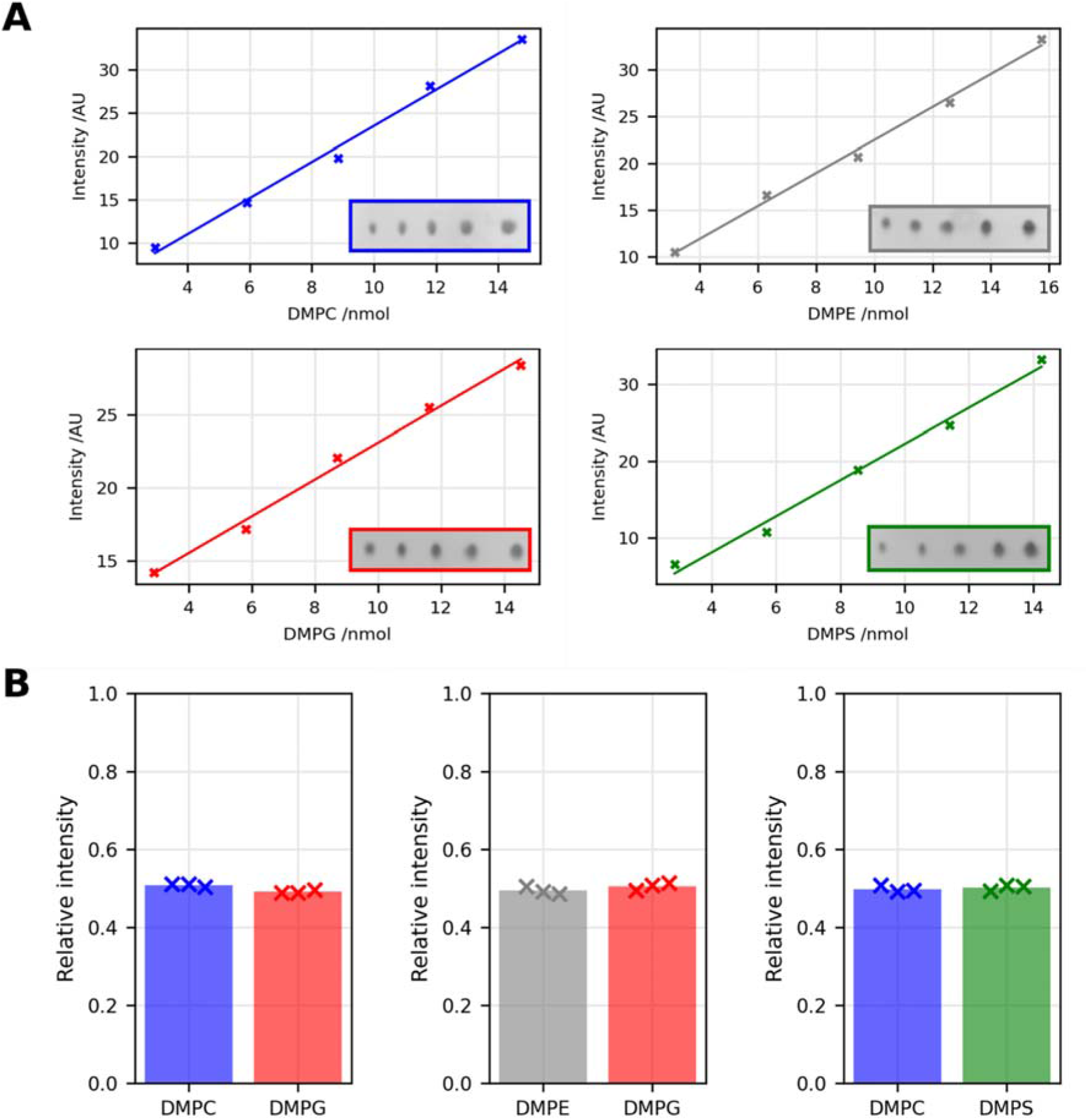
Thin layer chromatography (TLC) can be used to determine relative lipid ratios. **(A)** DMPC (blue), DMPG (red), DMPE (grey) and DMPS (green) lipid staining depth is directly and linearly proportional to amount of lipid loaded. One example from three replicates shown. **(B)** DMPC-DMPG, DMPE-DMPG and DMPS-DMPC lipids stain to equivalent depths for the same molar lipid amounts.

**Supplementary Figure 3:**
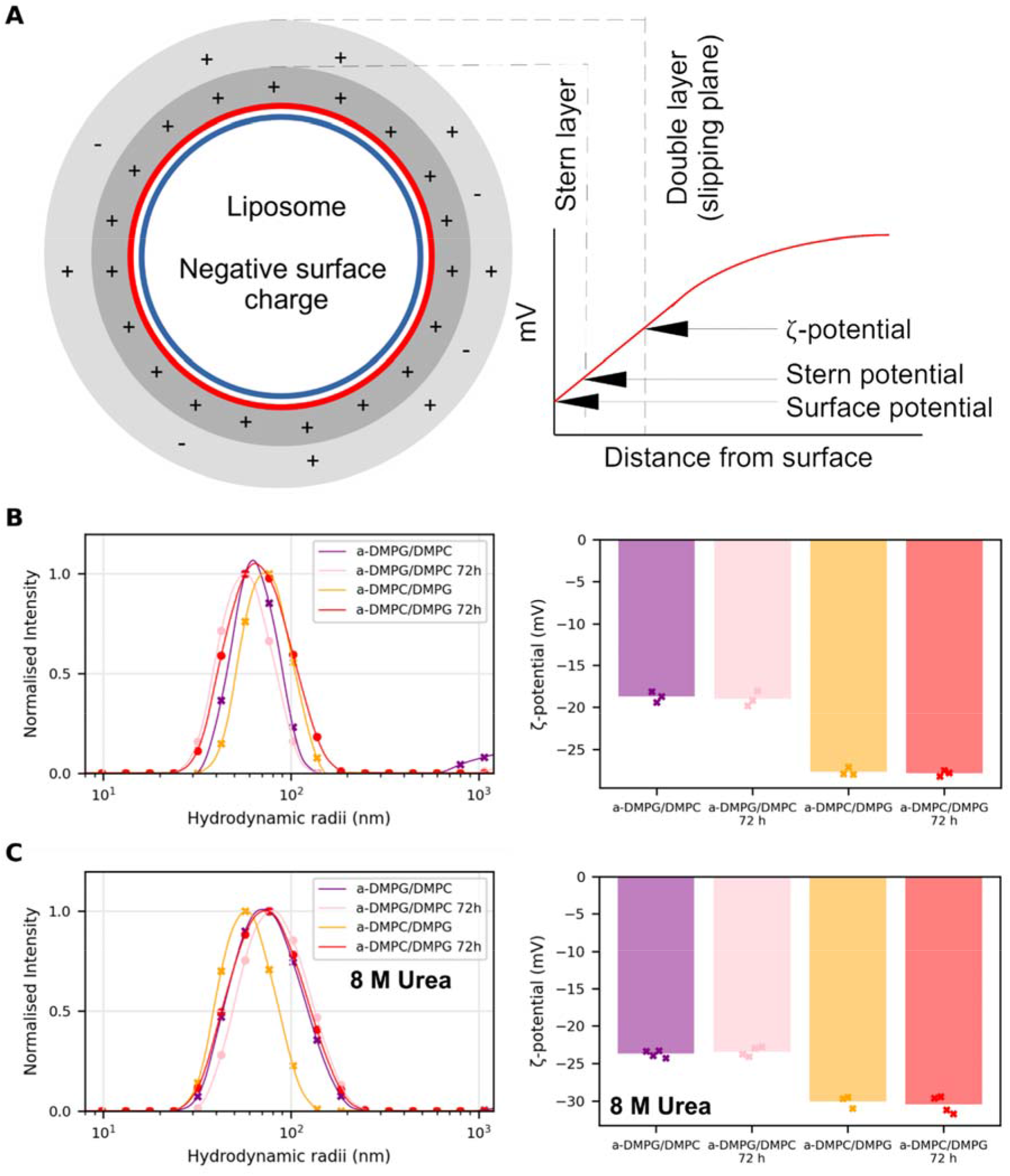
Liposome ζ-potential can be used to measure outer leaflet lipid composition. **(A)** Physical basis of the ζ-potential as the charge on the liposome slipping plane through solution (double layer). Liposome hydrodynamic radius and ζ-potential, and hence asymmetry, is stable **(B)** over at least 72 hours and **(C)** in 8 M urea. Note that asymmetry was determined without urea present.

**Supplementary Figure 4:**
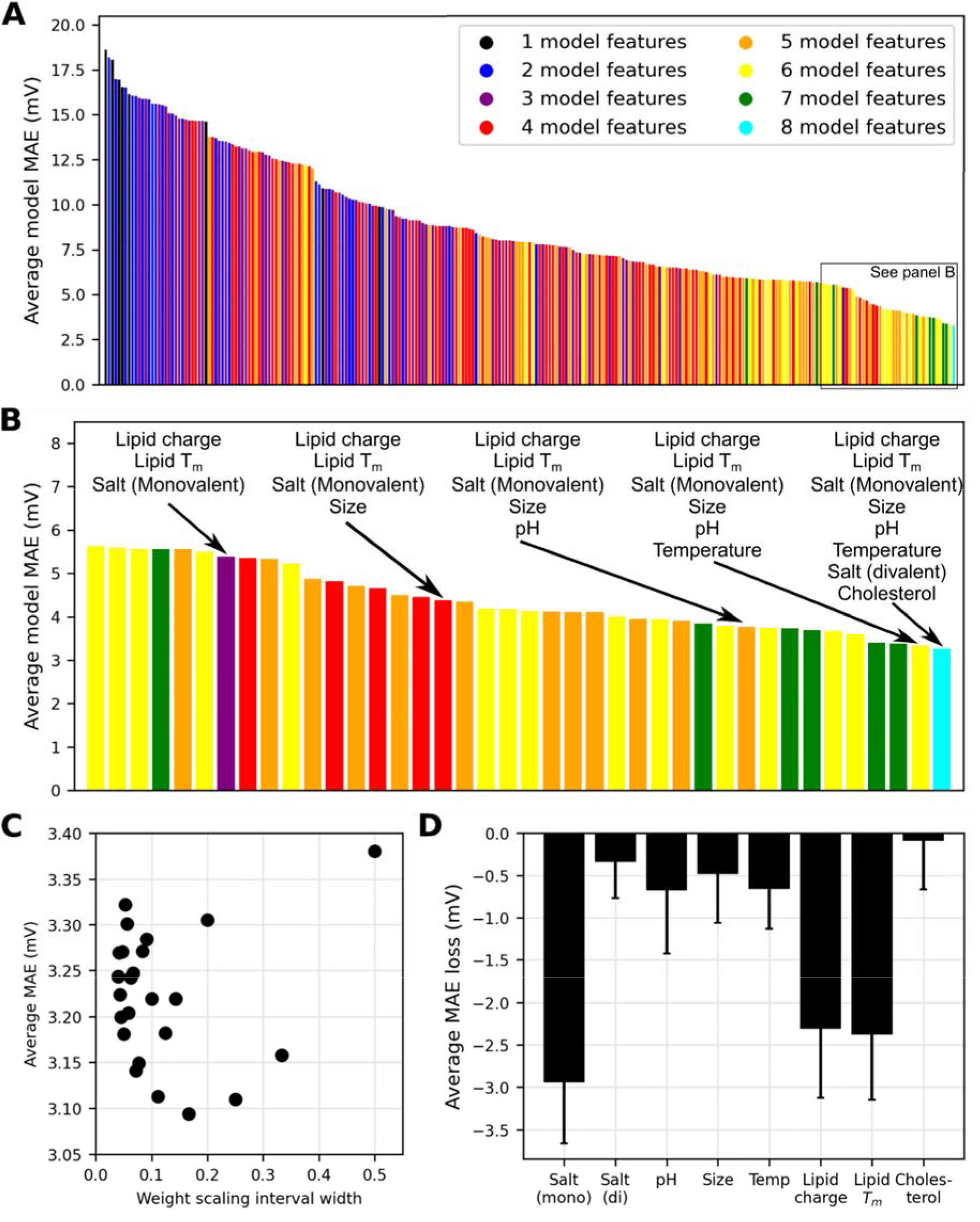
Understanding the ζ-potential prediction model. **(A)** The average MAE of models generated using all combinations of the eight dataset features (monovalent/divalent salt concentrations, pH, liposome size, temperature, cholesterol fraction and overall lipid charge/T_m_). **(B)** Expanded boxed region in (A). Considering the best model generated with reduced feature count indicates relative feature importance. **(C)** Effect of weighting the ζ-potential target values by their associated measurement error. Scaling intervals are centred on 0.5. An interval of 0.18 yields the best model improvement – i.e. all target values are weighted between 0.41 and 0.59 according to their error. **(D)** Average MAE loss from single parameter ablation from the final model. All averages MAEs from 50 trained models, error bars are standard deviation.

**Supplementary Figure 5:**
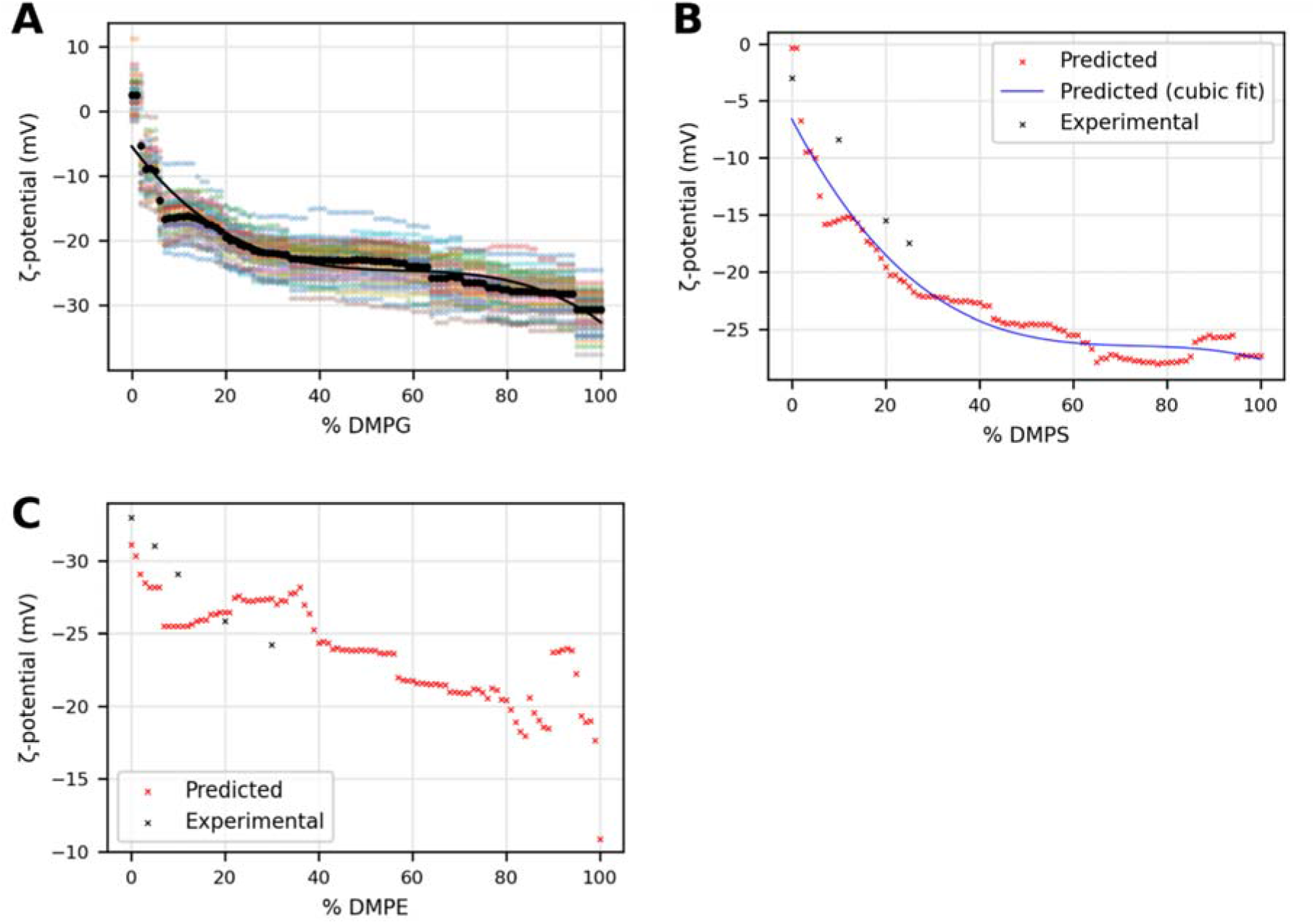
ζ-potential prediction model. **(A)**Individual predictions (coloured crosses) and their average (black circles) from the model ensemble for prediction in Figure 1H (DMPC-PG). The fitted cubic is also shown. **(B)** Average prediction for DMPS-PC and **(C)** DMPE-PG lipid mixes compared with the experimental data. Note that high-DMPE content LUVs (> 30-40% mol/mol) cannot be synthesised due to the strong negative curvature of DMPE.

**Supplementary Figure 6:**
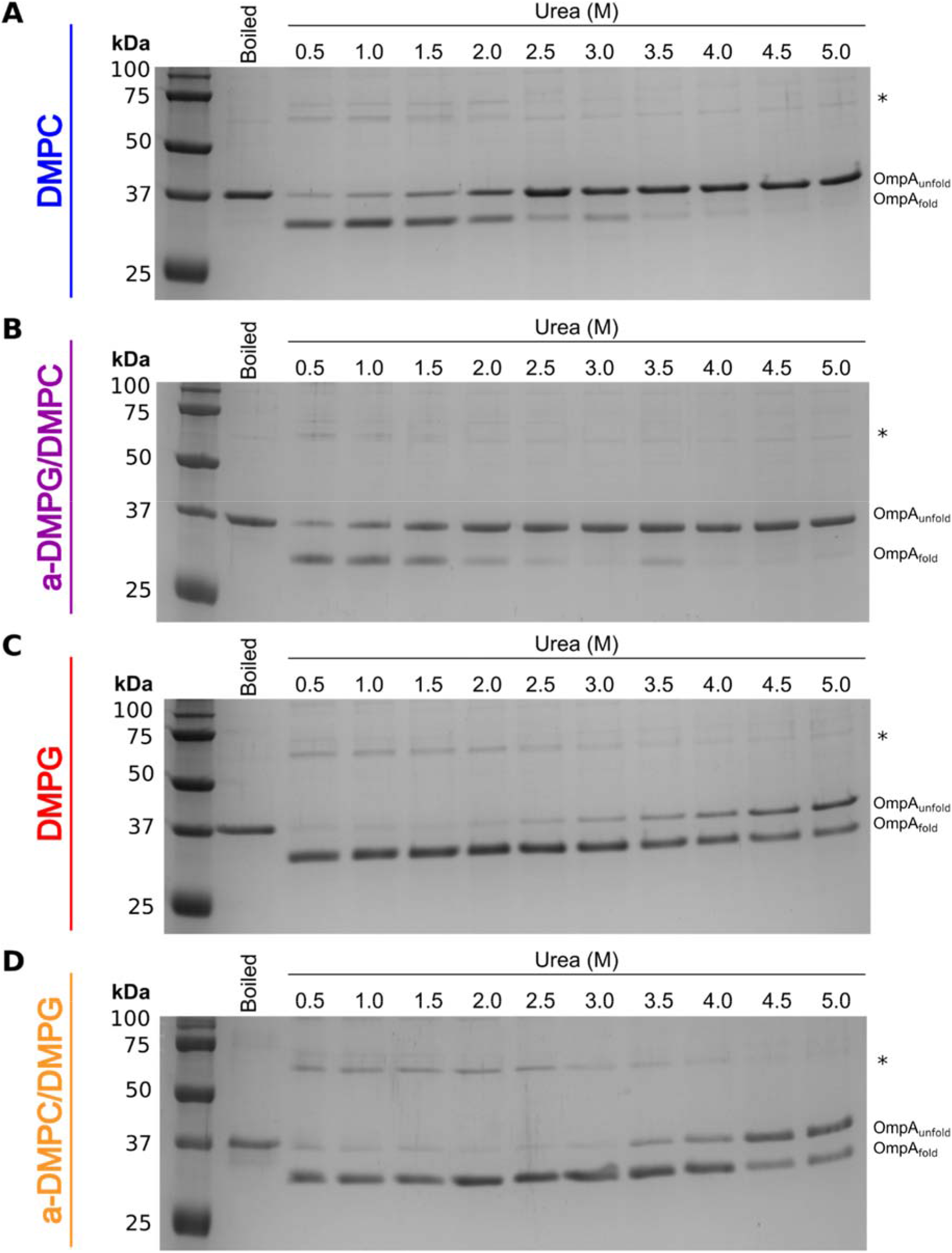
Example OmpA folding curves over 0.5 – 5 M urea in different liposome compositions. OmpA (initially unfolded in 8 M urea) was incubated with the relevant liposomes at each final concentration of urea indicated overnight at 30 °C. The proportion of folded protein was then analysed by cold SDS-PAGE (Methods) and the fraction folded determined from the folded and unfolded monomer bands only. Inclusion of higher order bands (indicated by *) in the densitometry analysis, or normalising against the boiled sample, showed minimal difference to final fraction folded (Methods).

**Supplementary Figure 7:**
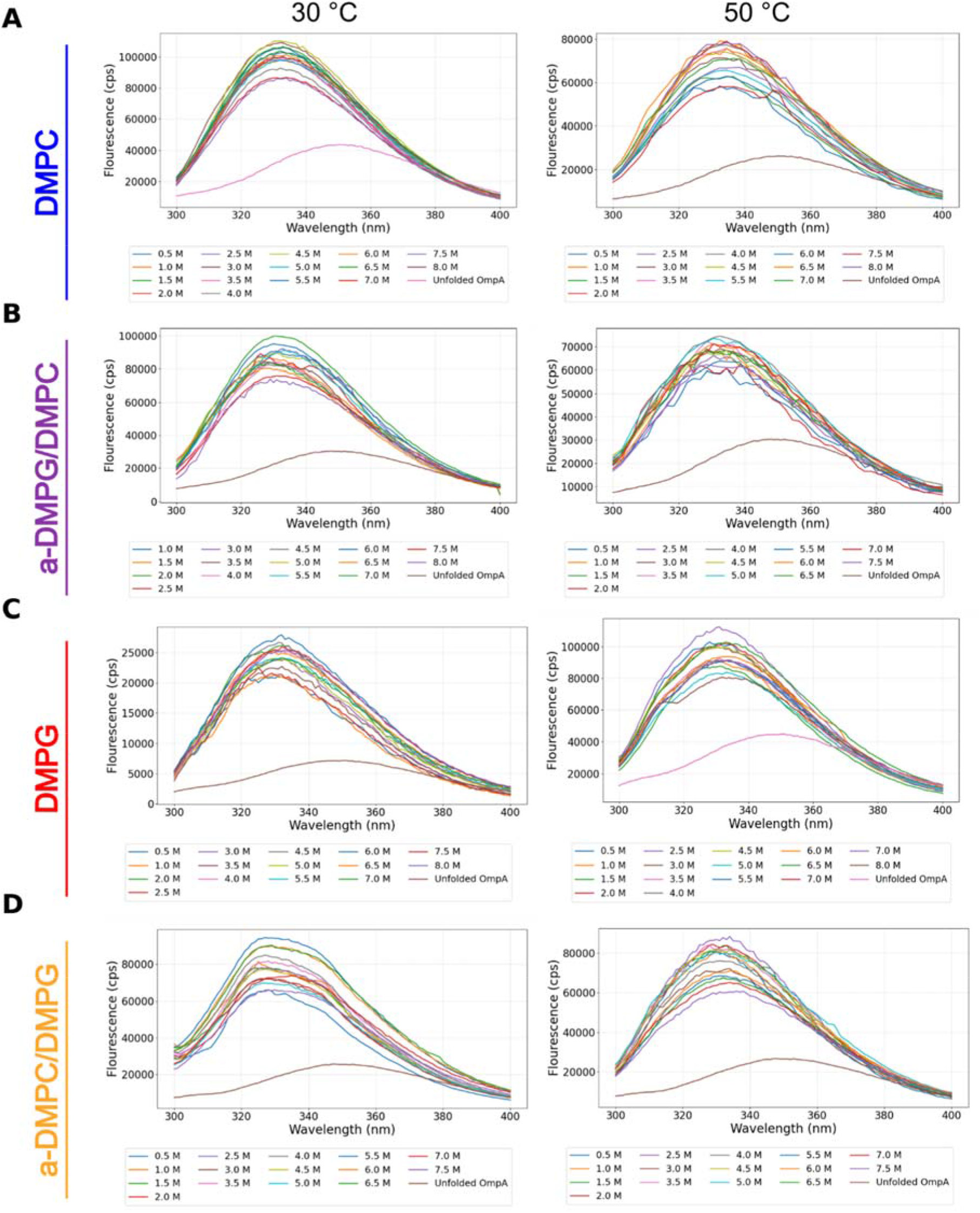
Urea-unfolding curves of OmpA measured by tryptophan fluorescence. (A-D) OmpA folded into LUVs of different lipid composition and organisation does not unfold following incubation at 30 °C or 50 °C overnight. The slight reduction in intensity indicates that the liposomes have started to aggregate during the overnight reaction. The spectrum of unfolded OmpA was determined in 7.5 M urea in the absence of lipid.

**Supplementary Figure 8:**
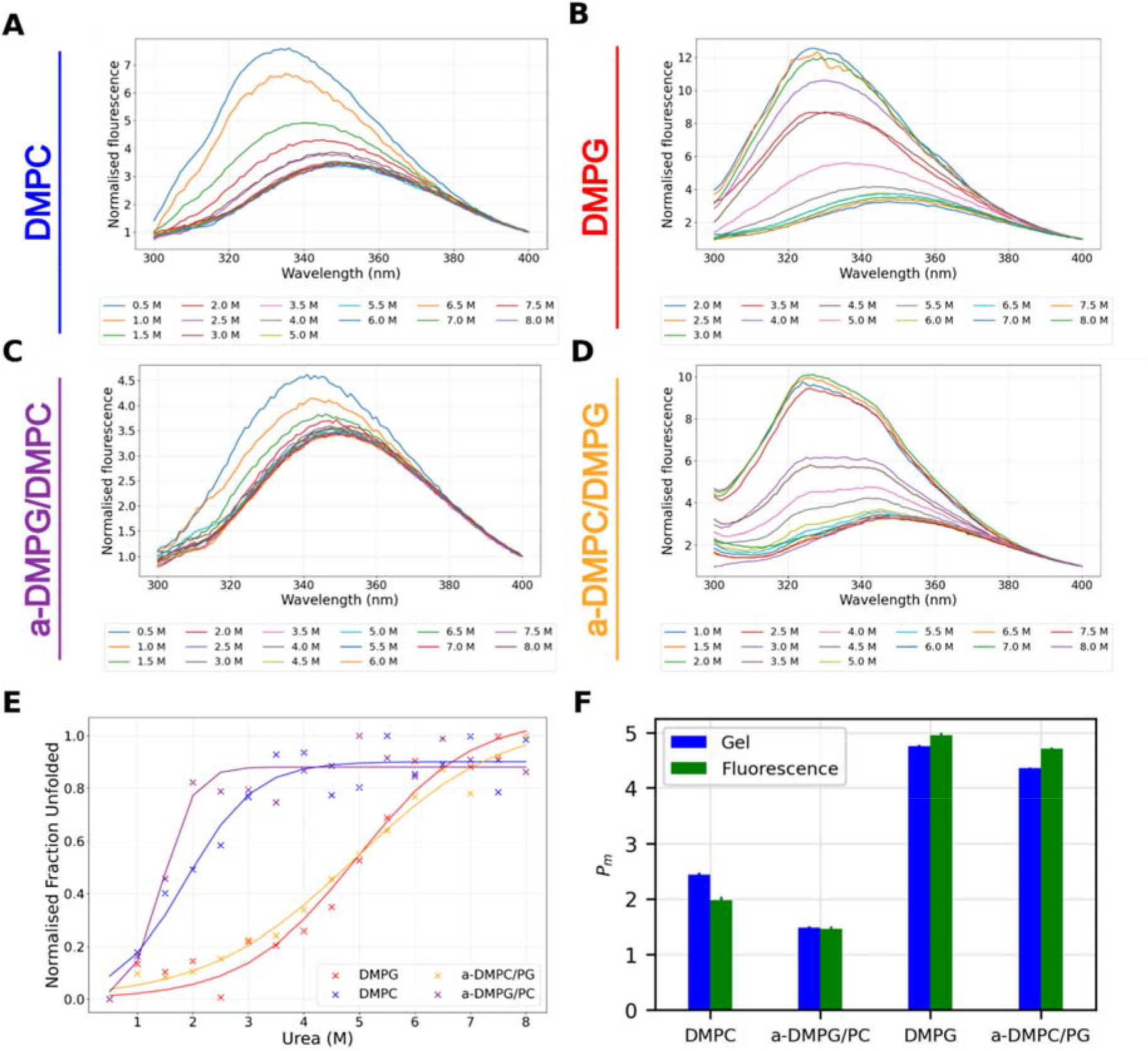
Urea-folding curves for OmpA in different liposome conditions measured by tryptophan fluorescence agree well with cold SDS-PAGE data. **(A-D)** Tryptophan fluorescence emission curves of Omp A in different liposome compositions and 0.5 – 8 M urea. **(E)** Extracting the 335/350 nm ratio of each spectrum and normalising to each liposome condition shows urea transition curves that **(F)** agree well with the cold SDS-PAGE data when compared by curve mid-points (differences less than 0.5 M urea, the increment size). Error bars indicate +/- goodness of fit (average difference between observed and fitted data).

**Supplementary Figure 9:**
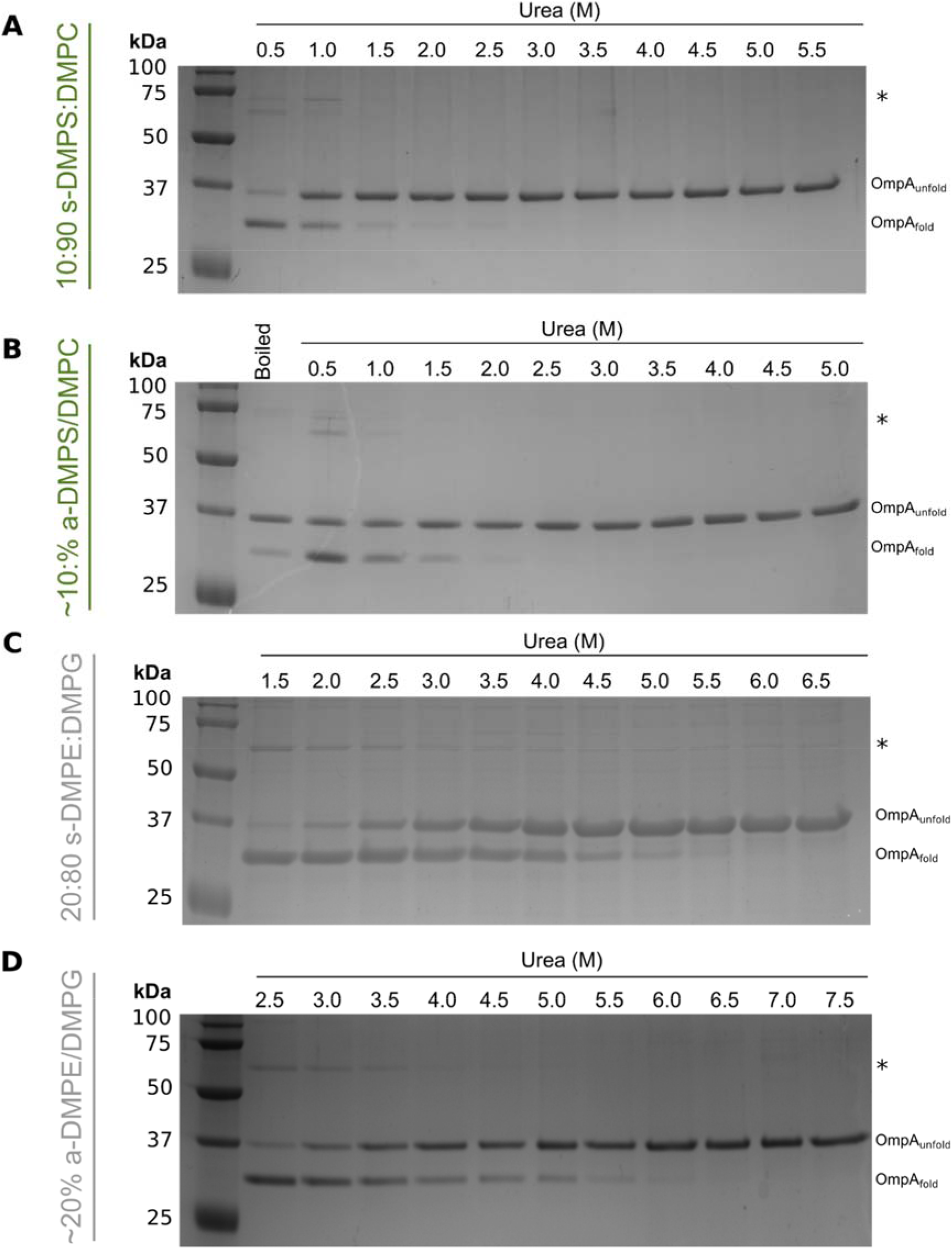
Example cold SDS PAGE gels for folding OmpA into DMPE and DMPS containing liposomes. **(A,B)**Example gels for OmpA folding into DMPS-DMPC symmetric and asymmetric liposomes at urea concentrations 0.5-5.5 M. **(C,D)** Example gels for OmpA folding into DMPE-DMPG symmetric and asymmetric liposomes at different urea concentrations. Note higher bands (*) not used to calculate midpoints, see Methods.

**Supplementary Figure 10:**
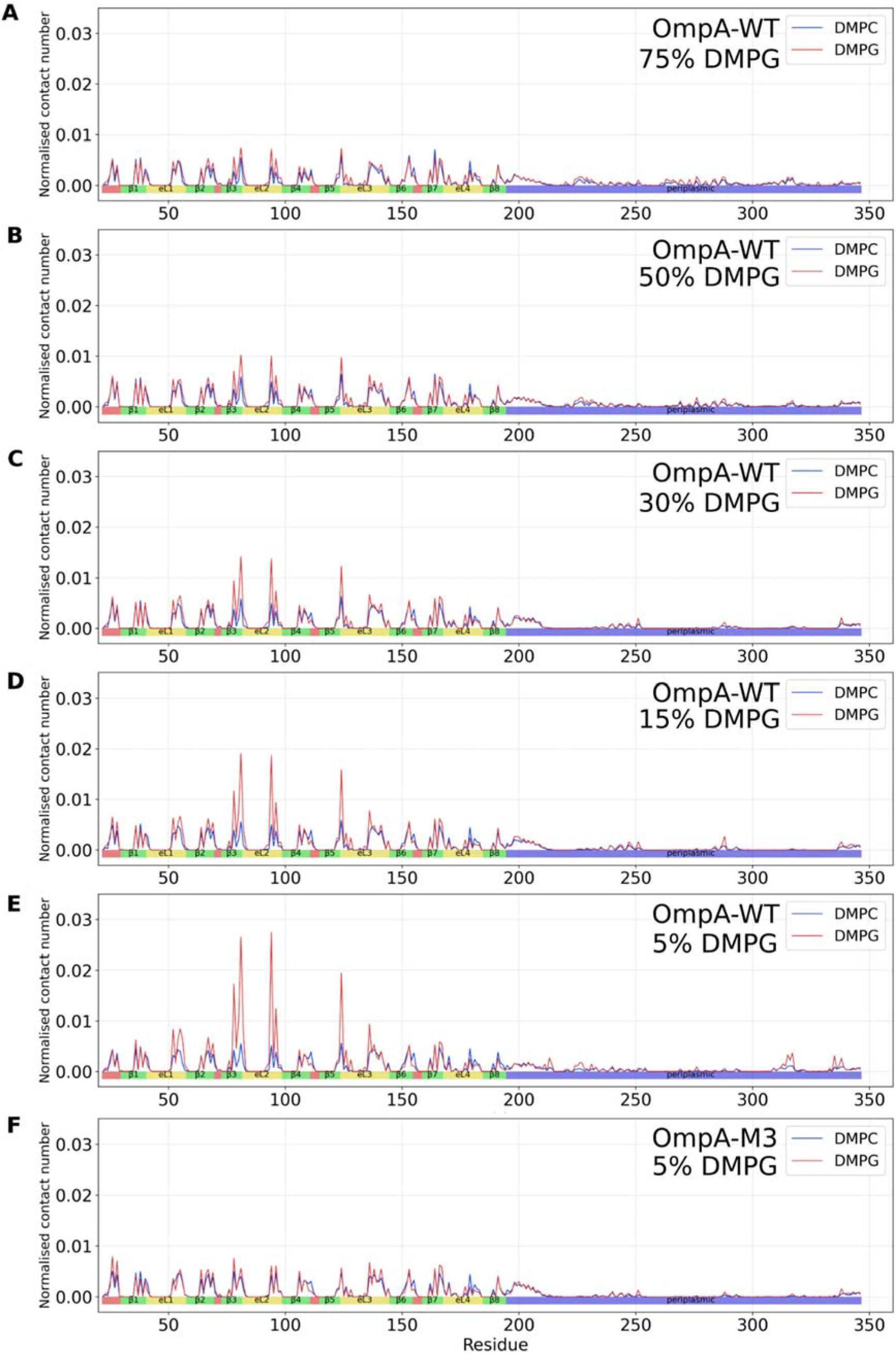
OmpA specifically interacts with DMPG in a titratable manner. **(A-E)**Normalised lipid-protein contacts between OmpA (WT) and DMPC or DMPG lipids with a total DMPG fraction of 75%, 50%, 30%, 15% or 5%. Note that the sequence of OmpA is shown below (strands (green), extracellular loops (yellow) intracellular turns (red), and includes its C- terminal water soluble domain (blue)). Peaks are more prominent with less DMPG due to a reduction of background signal. Normalised to lipid concentrations (Methods). **(F)** Normalised lipid-protein contacts between OmpA-M3 (R81S, K94S, R124S) and DMPC or DMPG lipids with total DMPG fraction of 5%. Comparing with WT OmpA (panel E), it is clear that the specific interactions with have been abrogated in the variant.

**Supplementary Figure 11:**
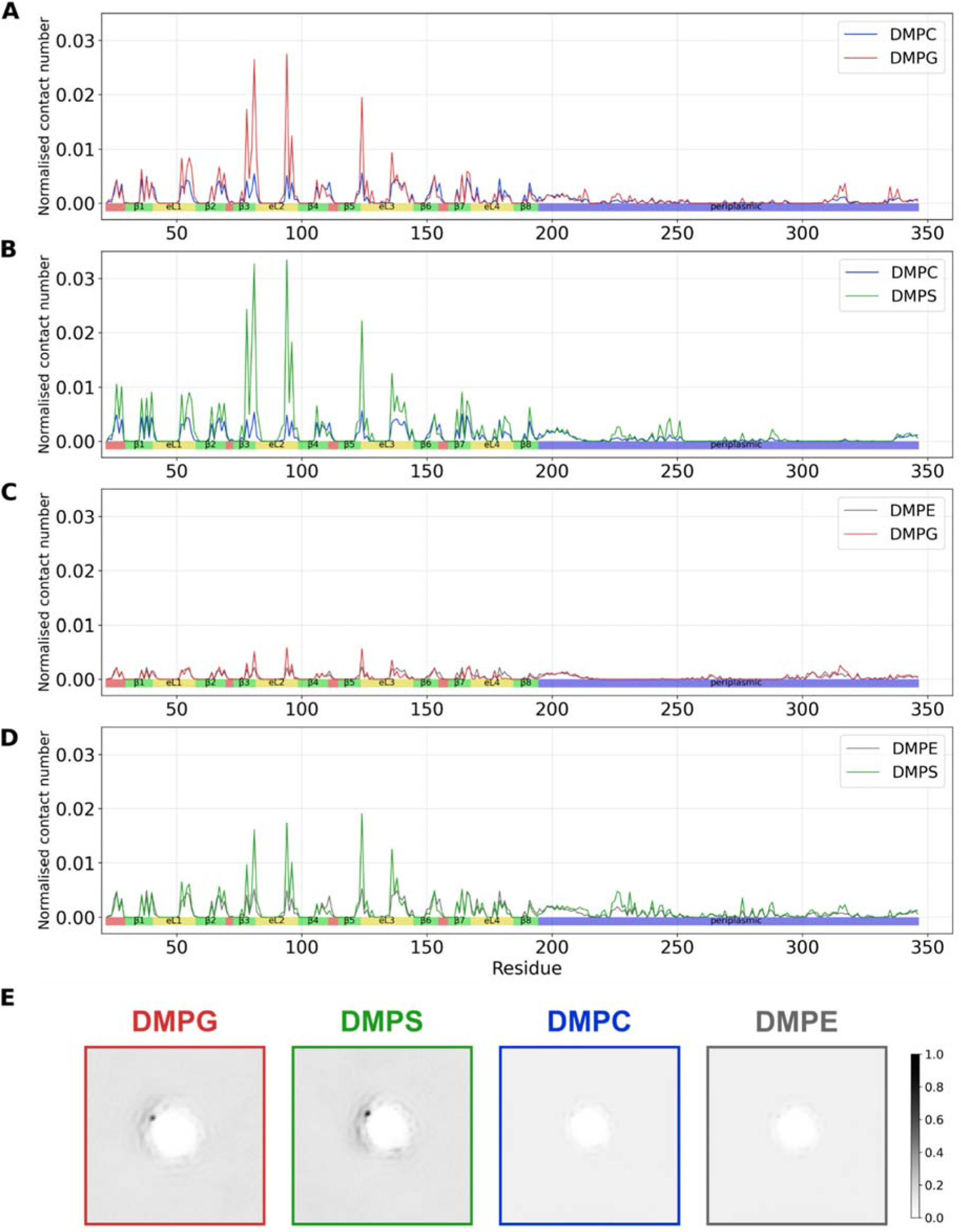
Normalised contact plots and densities for full-length OmpA interaction with DMPC, DMPG, DMPS and DMPE lipid mixes. Normalised contact plots for full length OmpA in (A) 5:95 s-DMPS:DMPG, (B) 5:95 s-DMPS:DMPC, (C) 5:95 s-DMPG:DMPE and (D) 5:95 s-DMPS:DMPE symmetric lipids. The sequence of OmpA is shown below (strands (green), extracellular loops (yellow) intracellular turns (red), and includes its C- terminal water soluble domain (blue)). (E) Lipid density plots for the phosphates of the respective lipids from both leaflets following centring and fitting of OmpA. Densities normalised to lipid concentrations. DMPC and DMPG calculated from 5:95 s-DMPC:DMPG, DMPS from 5:95 s-DMPS:DMPC and DMPE from 5:95 s-DMPG:DMPE systems.

**Supplementary Figure 12:**
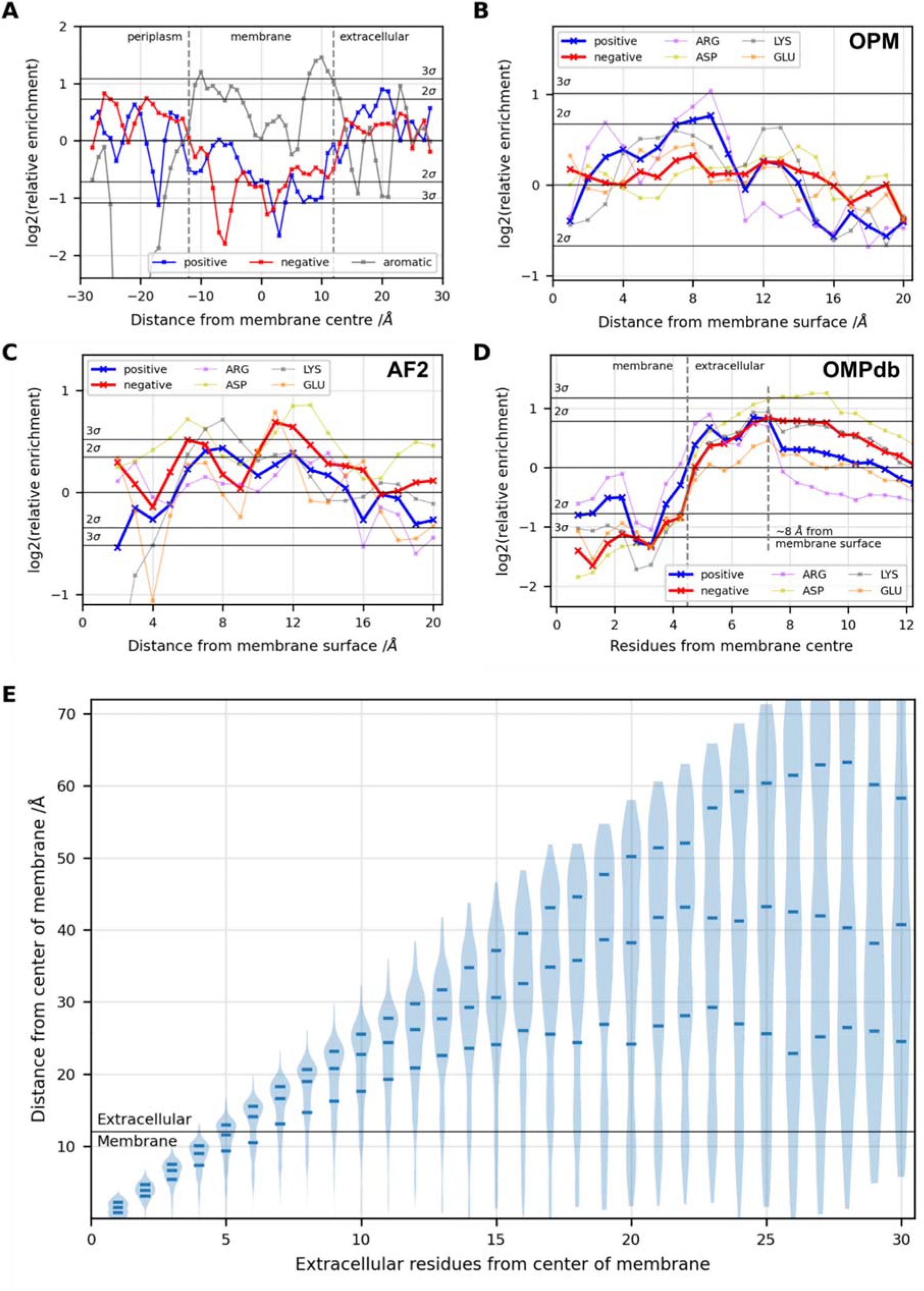
All charged residue enrichments relative to membrane proximity. Average enrichment of positively and negatively charged residues, and the underlying enrichment of arginine/lysine and aspartic/glutamic acid relative to random chance for the **(A-B)** OPM, **(C)** Alphafold2 (AF2) and **(D)** OMPdb datasets. **(E)** Violin plots indicating the average distance from the centre of the membrane for the Cα of residues in extracellular loops in the OPM dataset, showing the average and first and third quartile. Loops were truncated at their midpoints, and each part considered separately.

**Supplementary Figure 13:**
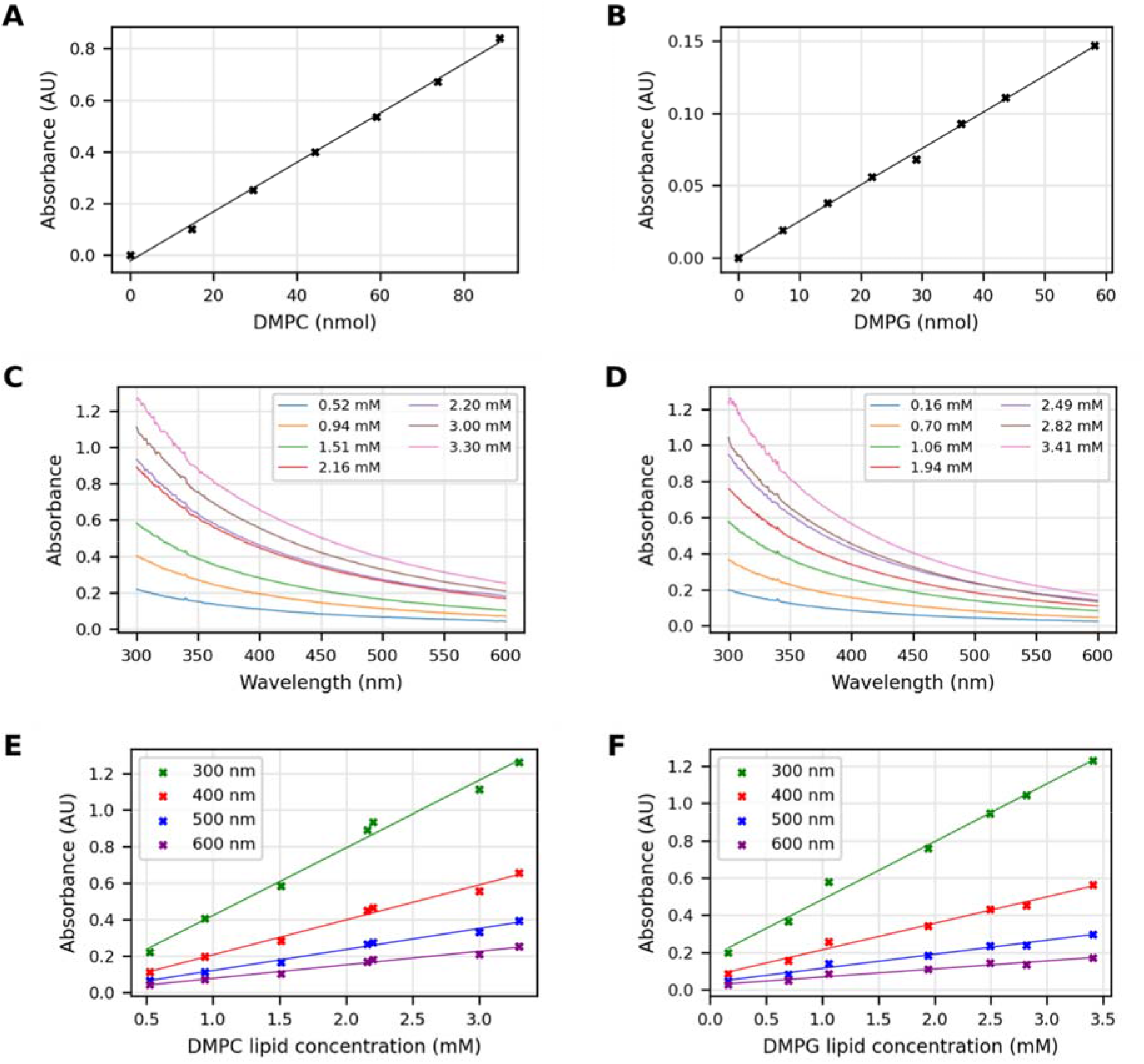
Determining liposome concentration. **(A,B)**Liposome absorbance calibration lines to absolute lipid amount, determined using the Stewart assay for DMPC and DMPG. Absorbance curves for **(C)** DMPC and **(D)** DMPG liposomes at different measured lipid concentration (measured using the Stewart assay). **(E,F)** The absorbance values at 300 nm, 400 nm, 500 nm and 600 nm of DMPC and DMPG LUVs, respectively, with fitted straight lines, to generate absorbance-liposome concentration calibration curves.

**Supplementary Figure 14:**
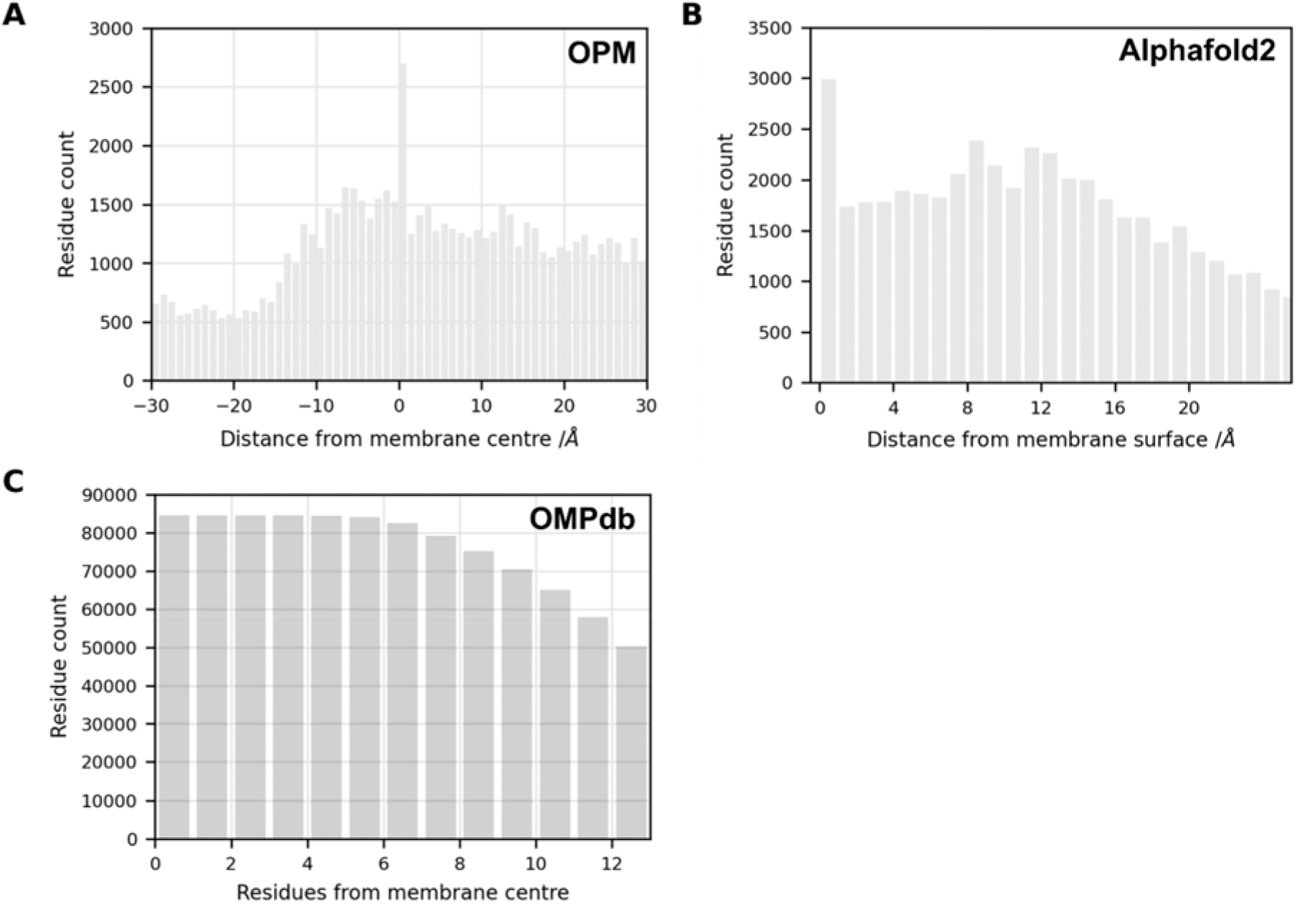
Number of residues per z-axis slab for (A) OPM, (B) Alphafold2 and (C) OMPdb residue enrichment analysis. Residue count used to calculate enrichments in each z-axis slab (per Å for OPM and Alphafold2 datasets, per residue for OMPdb).

**Supplementary Table 1:**
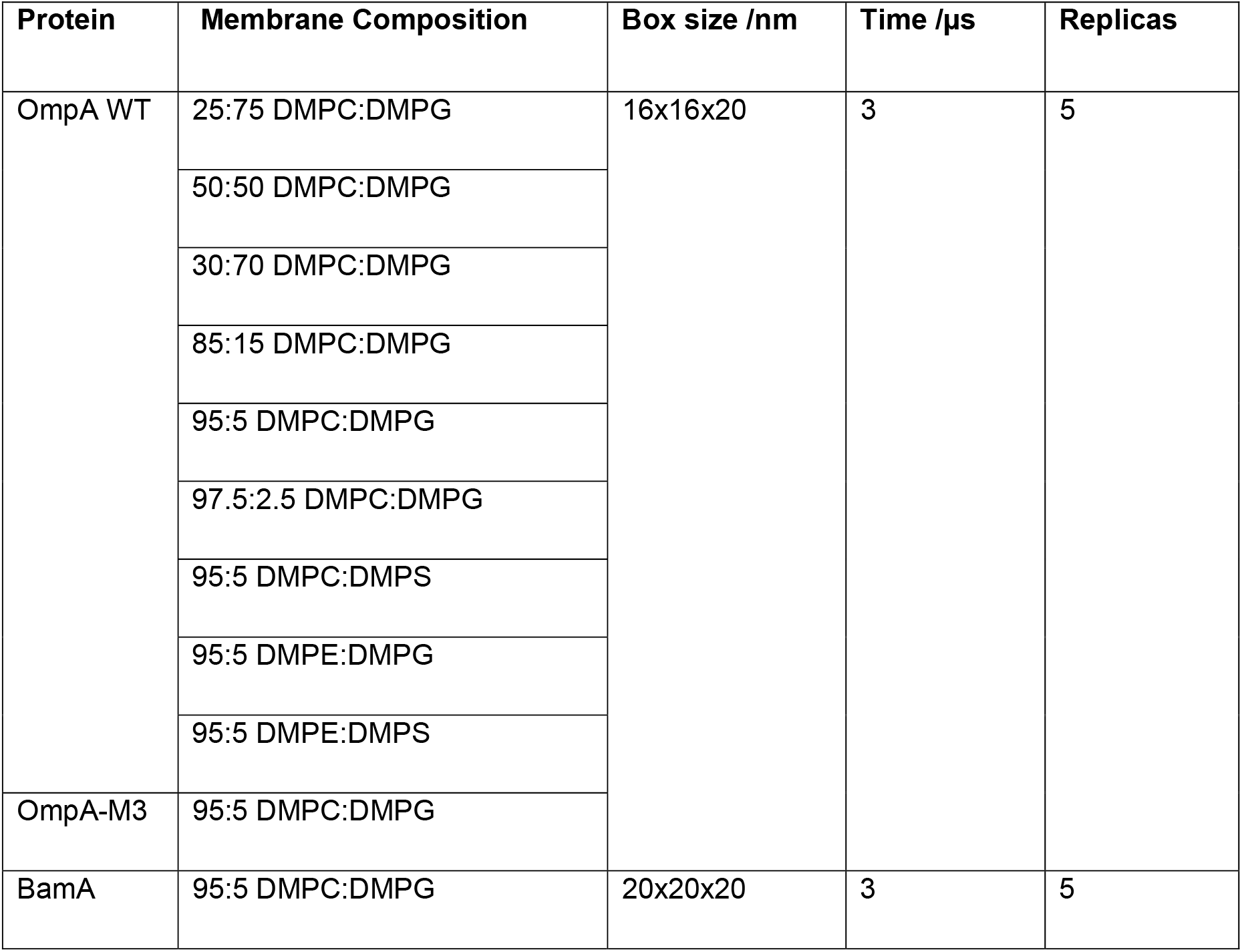
Summary of all simulations run in this study. Membrane composition fraction is by lipid number. OmpA-M3 is OmpA with R81S, K94S and R124S mutations.

| Protein | Membrane Composition | Box size /nm | Time /μs | Replicas |
| --- | --- | --- | --- | --- |
| OmpA WT | 25:75 DMPC:DMPG | 16x16x20 | 3 | 5 |
|  | 50:50 DMPC:DMPG |  |  |  |
|  | 30:70 DMPC:DMPG |  |  |  |
|  | 85:15 DMPC:DMPG |  |  |  |
|  | 95:5 DMPC:DMPG |  |  |  |
|  | 97.5:2.5 DMPC:DMPG |  |  |  |
|  | 95:5 DMPC:DMPS |  |  |  |
|  | 95:5 DMPE:DMPG |  |  |  |
|  | 95:5 DMPE:DMPS |  |  |  |
| OmpA-M3 | 95:5 DMPC:DMPG |  |  |  |
| BamA | 95:5 DMPC:DMPG | 20x20x20 | 3 | 5 |

## References

1. Rothman, J. E. & Lenard, J. Membrane asymmetry. Science 195, 743–753 (1977).

2. van Meer, G. Cellular lipidomics. EMBO J. 24, 3159–3165 (2005).

3. Situ, A. J. & Ulmer, T. S. Universal principles of membrane protein assembly, composition and evolution. PloS One 14, e0221372 (2019).

4. Dowhan, W., Vitrac, H. & Bogdanov, M. Lipid-Assisted Membrane Protein Folding and Topogenesis. Protein J. 38, 274–288 (2019).

5. Hong, H., Choi, H.-K. & Yoon, T.-Y. Untangling the complexity of membrane protein folding. Curr. Opin. Struct. Biol. 72, 237–247 (2022).

6. Fadeel, B. & Xue, D. The ins and outs of phospholipid asymmetry in the plasma membrane: roles in health and disease. Crit. Rev. Biochem. Mol. Biol. 44, 264–277 (2009).

7. Ikeda, M., Kihara, A. & Igarashi, Y. Lipid asymmetry of the eukaryotic plasma membrane: functions and related enzymes. Biol. Pharm. Bull. 29, 1542–1546 (2006).

8. Malinverni, J. C. & Silhavy, T. J. An ABC transport system that maintains lipid asymmetry in the gram-negative outer membrane. Proc. Natl. Acad. Sci. U. S. A. 106, 8009–8014 (2009).

9. Rivel, T., Ramseyer, C. & Yesylevskyy, S. The asymmetry of plasma membranes and their cholesterol content influence the uptake of cisplatin. Sci. Rep. 9, 5627 (2019).

10. Kurniawan, J., Ventrici de Souza, J. F., Dang, A. T., Liu, G. & Kuhl, T. L. Preparation and Characterization of Solid-Supported Lipid Bilayers Formed by Langmuir–Blodgett Deposition: A Tutorial. Langmuir 34, 15622–15639 (2018).

11. Michel, J. P., Wang, Y. X., Kiesel, I., Gerelli, Y. & Rosilio, V. Disruption of Asymmetric Lipid Bilayer Models Mimicking the Outer Membrane of Gram-Negative Bacteria by an Active Plasticin. Langmuir 33, 11028–11039 (2017).

12. Pautot, S., Frisken, B. J. & Weitz, D. A. Engineering asymmetric vesicles. Proc. Natl. Acad. Sci. 100, 10718–10721 (2003).

13. Paulowski, L. et al. The Beauty of Asymmetric Membranes: Reconstitution of the Outer Membrane of Gram-Negative Bacteria. Front. Cell Dev. Biol. 8, 586 (2020).

14. Enoki, T. A. & Feigenson, G. W. Asymmetric Bilayers by Hemifusion: Method and Leaflet Behaviors. Biophys. J. 117, 1037–1050 (2019).

15. Doktorova, M. et al. Preparation of asymmetric phospholipid vesicles for use as cell membrane models. Nat. Protoc. 13, 2086–2101 (2018).

16. Cheng, H.-T., Megha, null & London, E. Preparation and properties of asymmetric vesicles that mimic cell membranes: effect upon lipid raft formation and transmembrane helix orientation. J. Biol. Chem. 284, 6079–6092 (2009).

17. Markones, M. et al. Engineering Asymmetric Lipid Vesicles: Accurate and Convenient Control of the Outer Leaflet Lipid Composition. Langmuir ACS J. Surf. Colloids 34, 1999–2005 (2018).

18. St Clair, J. W., Kakuda, S. & London, E. Induction of Ordered Lipid Raft Domain Formation by Loss of Lipid Asymmetry. Biophys. J. 119, 483–492 (2020).

19. Lin, Q. & London, E. The influence of natural lipid asymmetry upon the conformation of a membrane-inserted protein (perfringolysin O). J. Biol. Chem. 289, 5467–5478 (2014).

20. Scott, H. L., Heberle, F. A., Katsaras, J. & Barrera, F. N. Phosphatidylserine Asymmetry Promotes the Membrane Insertion of a Transmembrane Helix. Biophys. J. 116, 1495–1506 (2019).

21. Verherstraeten, S. et al. Perfringolysin O: The Underrated Clostridium perfringens Toxin? Toxins 7, 1702–1721 (2015).

22. Reshetnyak, Y. K., Andreev, O. A., Segala, M., Markin, V. S. & Engelman, D. M. Energetics of peptide (pHLIP) binding to and folding across a lipid bilayer membrane. Proc. Natl. Acad. Sci. U. S. A. 105, 15340–15345 (2008).

23. Horne, J. E., Brockwell, D. J. & Radford, S. E. Role of the lipid bilayer in outer membrane protein folding in Gram-negative bacteria. J. Biol. Chem. 295, 10340–10367 (2020).

24. Schulz, G. E. beta-Barrel membrane proteins. Curr. Opin. Struct. Biol. 10, 443–447 (2000).

25. Schulz, G. E. The structure of bacterial outer membrane proteins. Biochim. Biophys. Acta 1565, 308–317 (2002).

26. Kleinschmidt, J. H. Folding of β-barrel membrane proteins in lipid bilayers - Unassisted and assisted folding and insertion. Biochim. Biophys. Acta 1848, 1927–1943 (2015).

27. Schiffrin, B. et al. Effects of Periplasmic Chaperones and Membrane Thickness on BamA-Catalyzed Outer-Membrane Protein Folding. J. Mol. Biol. 429, 3776–3792 (2017).

28. Burgess, N. K., Dao, T. P., Stanley, A. M. & Fleming, K. G. β-Barrel Proteins That Reside in the Escherichia coli Outer Membrane in Vivo Demonstrate Varied Folding Behavior in Vitro. J. Biol. Chem. 283, 26748–26758 (2008).

29. Danoff, E. J. & Fleming, K. G. Membrane defects accelerate outer membrane β-barrel protein folding. Biochemistry 54, 97–99 (2015).

30. Patel, G. J. & Kleinschmidt, J. H. The lipid bilayer-inserted membrane protein BamA of Escherichia coli facilitates insertion and folding of outer membrane protein A from its complex with Skp. Biochemistry 52, 3974–3986 (2013).

31. Gessmann, D. et al. Outer membrane β-barrel protein folding is physically controlled by periplasmic lipid head groups and BamA. Proc. Natl. Acad. Sci. U. S. A. 111, 5878–5883 (2014).

32. Tiwari, P. B. & Mahalakshmi, R. Interplay of protein primary sequence, lipid membrane, and chaperone in β-barrel assembly. Protein Sci. Publ. Protein Soc. 30, 624–637 (2021).

33. Peterson, J. H., Plummer, A. M., Fleming, K. G. & Bernstein, H. D. Selective pressure for rapid membrane integration constrains the sequence of bacterial outer membrane proteins. Mol. Microbiol. 106, 777–792 (2017).

34. Schiffrin, B., Brockwell, D. J. & Radford, S. E. Outer membrane protein folding from an energy landscape perspective. BMC Biol. 15, 123 (2017).

35. Zhou, H.-X. & Pang, X. Electrostatic Interactions in Protein Structure, Folding, Binding, and Condensation. Chem. Rev. 118, 1691 (2018).

36. van Klompenburg, W., Nilsson, I., von Heijne, G. & de Kruijff, B. Anionic phospholipids are determinants of membrane protein topology. EMBO J. 16, 4261–4266 (1997).

37. Nilsson, J., Persson, B. & von Heijne, G. Comparative analysis of amino acid distributions in integral membrane proteins from 107 genomes. Proteins 60, 606–616 (2005).

38. Jackups, R. & Liang, J. Interstrand pairing patterns in beta-barrel membrane proteins: the positive outside rule, aromatic rescue, and strand registration prediction. J. Mol. Biol. 354, 979–993 (2005).

39. Slusky, J. S. G., Dunbrack, R. L. Jr. Charge asymmetry in the proteins of the outer membrane. Bioinformatics 29, 2122 (2013).

40. Koebnik, R. Structural and functional roles of the surface-exposed loops of the beta-barrel membrane protein OmpA from Escherichia coli. J. Bacteriol. 181, 3688–3694 (1999).

41. Vasan, A. K. et al. Role of internal loop dynamics in antibiotic permeability of outer membrane porins. Proc. Natl. Acad. Sci. U. S. A. 119, e2117009119 (2022).

42. Smith, M. C., Crist, R. M., Clogston, J. D. & McNeil, S. E. Zeta potential: a case study of cationic, anionic, and neutral liposomes. Anal. Bioanal. Chem. 409, 5779–5787 (2017).

43. Soema, P. C., Willems, G.-J., Jiskoot, W., Amorij, J.-P. & Kersten, G. F. Predicting the influence of liposomal lipid composition on liposome size, zeta potential and liposome-induced dendritic cell maturation using a design of experiments approach. Eur. J. Pharm. Biopharm. Off. J. Arbeitsgemeinschaft Pharm. Verfahrenstechnik EV 94, 427–435 (2015).

44. Sęk, A., Perczyk, P., Wydro, P., Gruszecki, W. I. & Szcześ, A. Effect of trace amounts of ionic surfactants on the zeta potential of DPPC liposomes. Chem. Phys. Lipids 235, 105059 (2021).

45. Svirina, A. & Terterov, I. Electrostatic effects in saturation of membrane binding of cationic cell penetrating peptide. Eur. Biophys. J. EBJ 50, 15–23 (2021).

46. Makino, K. et al. Temperature and ionic strength-induced conformational changes in the lipid head group region of liposomes as suggested by zeta potential data. Biophys. Chem. 41, 175–183 (1991).

47. Silvander, M., Hansson, P. & Edwards, K. Liposomal surface potential and bilayer packing as affected by PEG-lipid inclusion. Langmuir 16, 3696–3702 (2000).

48. Tatulian, S. A. Effect of lipid phase transition on the binding of anions to dimyristoylphosphatidylcholine liposomes. Biochim. Biophys. Acta 736, 189–195 (1983).

49. Lairion, F. & Disalvo, E. A. Effect of Dipole Potential Variations on the Surface Charge Potential of Lipid Membranes. J. Phys. Chem. B 113, 1607–1614 (2009).

50. Le, Q.-C., Ropers, M.-H., Terrisse, H. & Humbert, B. Interactions between phospholipids and titanium dioxide particles. Colloids Surf. B Biointerfaces 123, 150–157 (2014).

51. Luzardo, M., Peltzer, G. & Disalvo, E. Surface potential of lipid interfaces formed by mixtures of phosphatidylcholine of different chain lengths. (1998) doi:10.1021/LA971273J.

52. Markones, M. et al. Stairway to Asymmetry: Five Steps to Lipid-Asymmetric Proteoliposomes. Biophys. J. 118, 294–302 (2020).

53. Fatouros, D. G. & Antimisiaris, S. G. Effect of amphiphilic drugs on the stability and zeta-potential of their liposome formulations: a study with prednisolone, diazepam, and griseofulvin. J. Colloid Interface Sci. 251, 271–277 (2002).

54. Morini, M. A. et al. Influence of temperature, anions and size distribution on the zeta potential of DMPC, DPPC and DMPE lipid vesicles. Colloids Surf. B Biointerfaces 131, 54–58 (2015).

55. Andersen, K. K., Wang, H. & Otzen, D. E. A kinetic analysis of the folding and unfolding of OmpA in urea and guanidinium chloride: single and parallel pathways. Biochemistry 51, 8371–8383 (2012).

56. Bulieris, P. V., Behrens, S., Holst, O. & Kleinschmidt, J. H. Folding and insertion of the outer membrane protein OmpA is assisted by the chaperone Skp and by lipopolysaccharide. J. Biol. Chem. 278, 9092–9099 (2003).

57. Danoff, E. J. & Fleming, K. G. The soluble, periplasmic domain of OmpA folds as an independent unit and displays chaperone activity by reducing the self-association propensity of the unfolded OmpA transmembrane β-barrel. Biophys. Chem. 159, 194–204 (2011).

58. Surrey, T. & Jähnig, F. Refolding and oriented insertion of a membrane protein into a lipid bilayer. Proc. Natl. Acad. Sci. U. S. A. 89, 7457–7461 (1992).

59. Pocanschi, C. L., Patel, G. J., Marsh, D. & Kleinschmidt, J. H. Curvature Elasticity and Refolding of OmpA in Large Unilamellar Vesicles. Biophys. J. 91, L75 (2006).

60. Caffrey, M. & Hogan, J. LIPIDAT: a database of lipid phase transition temperatures and enthalpy changes. DMPC data subset analysis. Chem. Phys. Lipids 61, 1–109 (1992).

61. White, P. et al. The role of membrane destabilisation and protein dynamics in BAM catalysed OMP folding. Nat. Commun. 12, 4174 (2021).

62. Jumper, J. et al. Highly accurate protein structure prediction with AlphaFold. Nature 596, 583–589 (2021).

63. Varadi, M. et al. AlphaFold Protein Structure Database: massively expanding the structural coverage of protein-sequence space with high-accuracy models. Nucleic Acids Res. 50, D439–D444 (2022).

64. Tsirigos, K. D., Bagos, P. G. & Hamodrakas, S. J. OMPdb: a database of {beta}-barrel outer membrane proteins from Gram-negative bacteria. Nucleic Acids Res. 39, D324–331 (2011).

65. Lundstedt, E., Kahne, D. & Ruiz, N. Assembly and maintenance of lipids at the bacterial outer membrane. Chem. Rev. 121, 5098–5123 (2021).

66. Stewart, J. C. Colorimetric determination of phospholipids with ammonium ferrothiocyanate. Anal. Biochem. 104, 10–14 (1980).

67. Chen, T. & Guestrin, C. XGBoost: A Scalable Tree Boosting System. in Proceedings of the 22nd ACM SIGKDD International Conference on Knowledge Discovery and Data Mining 785–794 (Association for Computing Machinery, 2016). doi:10.1145/2939672.2939785.

68. Danoff, E. J. & Fleming, K. G. Novel Kinetic Intermediates Populated along the Folding Pathway of the Transmembrane β-Barrel OmpA. Biochemistry 56, 47–60 (2017).

69. Van Der Spoel, D., et al. GROMACS: fast, flexible, and free. J. Comput. Chem. 26, 1701–1718 (2005).

70. de Jong, D. H. et al. Improved Parameters for the Martini Coarse-Grained Protein Force Field. J. Chem. Theory Comput. 9, 687–697 (2013).

71. Marrink, S. J., Risselada, H. J., Yefimov, S., Tieleman, D. P. & de Vries, A. H. The MARTINI Force Field: Coarse Grained Model for Biomolecular Simulations. J. Phys. Chem. B 111, 7812–7824 (2007).

72. Wassenaar, T. A., Ingólfsson, H. I., Böckmann, R. A., Tieleman, D. P. & Marrink, S. J. Computational Lipidomics with insane: A Versatile Tool for Generating Custom Membranes for Molecular Simulations. J. Chem. Theory Comput. 11, 2144–2155 (2015).

73. Parrinello, M. & Rahman, A. Polymorphic transitions in single crystals: A new molecular dynamics method. J. Appl. Phys. 52, 7182–7190 (1981).

74. Bussi, G., Donadio, D. & Parrinello, M. Canonical sampling through velocity rescaling. J. Chem. Phys. 126, 014101 (2007).

75. Hess, B., Bekker, H., Berendsen, H. J. C. & Fraaije, J. G. E. M. LINCS: A linear constraint solver for molecular simulations. J. Comput. Chem. 18, 1463–1472 (1997).

76. Song, W. et al. PyLipID: A Python Package for Analysis of Protein–Lipid Interactions from Molecular Dynamics Simulations. J. Chem. Theory Comput. 18, 1188–1201 (2022).

77. Lomize, M. A., Lomize, A. L., Pogozheva, I. D. & Mosberg, H. I. OPM: orientations of proteins in membranes database. Bioinforma. Oxf. Engl. 22, 623–625 (2006).

78. Fu, L., Niu, B., Zhu, Z., Wu, S. & Li, W. CD-HIT: accelerated for clustering the next-generation sequencing data. Bioinformatics 28, 3150–3152 (2012).

79. Almagro Armenteros, J. J., et al. SignalP 5.0 improves signal peptide predictions using deep neural networks. Nat. Biotechnol. 37, 420–423 (2019).

